# The juvenile hormone receptor Methoprene-tolerant is involved in the sterilizing effect of pyriproxyfen on adult *Aedes aegypti* mosquitoes

**DOI:** 10.1101/2020.05.31.126508

**Authors:** Tahmina Hossain Ahmed, T. Randolph Saunders, Donald Mullins, Mohammad Zillur Rahman, Jinsong Zhu

**Author notes:** Corresponding author (JZ).

## Abstract

Exposure of adult mosquitoes to pyriproxyfen (PPF), an analog of insect juvenile hormone (JH), has shown promise to effectively sterilize female mosquitoes. However, the underlying mechanisms of the PPF-induced decrease in mosquito fecundity are largely unknown. We performed a comprehensive study to dissect the mode of PPF action in *Aedes aegypti* mosquitoes. Exposure to PPF prompted the overgrowth of primary follicles in sugar-fed *Ae. aegypti* females but blocked the development of primary follicles at Christopher’s Stage III after blood feeding. Secondary follicles were precociously activated in PPF-treated mosquitoes. Moreover, PPF substantially altered the expression of many genes that are essential for mosquito physiology and oocyte development in the fat body and ovary. In particular, many metabolic genes were differentially expressed in response to PPF treatment, thereby affecting the mobilization and utilization of energy reserves. Furthermore, PPF treatment on the previtellogenic female adults considerably modified mosquito responses to JH and 20-hydroxyecdysone (20E), two major hormones that govern mosquito reproduction. Krüppel homolog 1, a JH-inducible transcriptional regulator, showed consistently elevated expression after PPF exposure. Conversely, PPF upregulated the expression of several key players of the 20E regulatory cascades, including *HR3* and *E75A*, in the previtellogenic stage. After blood-feeding, the expression of these 20E response genes was significantly weaker in PPF-treated mosquitoes than the solvent-treated control groups. RNAi-mediated knockdown of the Methoprene-tolerant (Met) protein, the JH receptor, partially rescued the impaired follicular development after PPF exposure and substantially increased the hatching of the eggs produced by PPF-treated female mosquitoes. Thus, the results suggested that PPF relied on Met to exert its sterilizing effects on female mosquitoes. In summary, this study finds that PPF exposure disturbs normal hormonal responses and metabolism in *Ae. aegypti*, shedding light on the molecular targets and the downstream signaling pathways activated by PPF.

**Author summary:** *Aedes aegypti* mosquitoes are responsible for the transmission of dengue, yellow fever, chikungunya, and Zika fever. Insecticides are widely used as the primary tool in the prevention and control of these infectious diseases. In light of the rapid increase of insecticide resistance in mosquito populations, there is an urgent need to find new classes of insecticides with a different mode of action. Here we found that pyriproxyfen, an analog of insect juvenile hormone (JH), had a large impact on the oocyte development, both before and after blood feeding, in female mosquitoes. Pyriproxyfen disturbed normal hormonal responses and caused metabolic shifting in female adults. These actions appear to collectively impair oocyte development and substantially reduce viable progenies of female mosquitoes. Besides, we demonstrated the involvement of the JH receptor Met in pyriproxyfen-induced female sterilization. This study significantly advances our understanding of mosquito reproductive biology and the molecular basis of pyriproxyfen action, which are invaluable for the development of new mosquito control strategies.

## Introduction

The yellow fever mosquito, *Aedes aegypti,* is a major vector of several globally important human arboviral diseases including zika, dengue, chikungunya, and yellow fever [1]. Currently, there is still a lack of effective vaccines and specific antiviral therapies for most of these mosquito-borne diseases. Pyrethroids are the most common insecticides used to reduce mosquito populations [2]. They comprise ~40% of all insecticides used annually and are the only insecticide recommended by the World Health Organization for the treatment of bed nets [3]. However, insecticide resistance evolves readily in mosquito populations and is jeopardizing the effectiveness of several commonly used insecticides including pyrethroids [4–7]. Therefore, new mosquito insecticides with different modes of action are urgently needed.

Juvenile hormone (JH), due to its exclusive presence in arthropods, has attracted a lot of attention in the development of environment-friendly insecticides [8]. Pyriproxyfen (PPF), a synthetic JH analog, is used as a larvicide that interferes with the development of mosquito larvae preventing the emergence of adults [9]. Moreover, sublethal exposure to PPF at larval stages has deleterious effects on egg development and fertility of surviving adult mosquitoes [10,11]. Direct topical application of PPF to adult female mosquitoes considerably reduces egg production and decreases the hatching rate of oviposited eggs [12–16]. PPF-treated bed nets have shown promise in suppressing mosquito reproduction and reducing mosquito populations under semi-field conditions [14]. Several studies have reported that JH analogs, including methoprene and PPF, delay or stall the follicular development in adult female *Ae. aegypti* and *Anopheles gambiae* mosquitoes [12,17,18]. However, little is known about how PPF hinders the development of ovarian follicles or whether follicular development is arrested at a particular stage.

Oogenesis in adult female mosquitoes is controlled by JH and the ecdysteroid hormone, 20-hydroxyecdysone (20E) [19,20]. Female adults in most mosquito species require a blood meal from vertebrate hosts for their egg maturation [21]. Separated by blood-feeding, the first gonadotrophic cycle can be divided into the previtellogenic phase (before a blood meal) and the vitellogenic phase (after blood ingestion). In the first three days after eclosion, JH prompts the growth of primary follicles to reach a length of about 100 μm [22]. The follicles then enter an arrested stage of development awaiting a blood meal. Synthesis of 20E is triggered by blood-feeding, which also leads to a rapid decrease in JH levels. 20E controls the synthesis of yolk protein precursors in the fat body and the progression of oogenesis in the ovary. At about 30 h post blood meal (PBM), 20E drops back to basal levels, signaling the end of vitellogenic synthesis. Egg maturation is completed by 72 h PBM and female mosquitoes are ready for oviposition. Conversely, JH titers rise again starting at 48 h PBM, initiating previtellogenic growth of secondary follicles and priming mosquitoes for the second gonotrophic cycle.

Adult female mosquitoes typically obtain nutrients for survival and reproduction from three sources: larval-derived teneral reserves, nectar or other plant juices, and blood meals. Metabolic processes in mosquitoes, which constantly react to nutritional changes to meet various energy demands, are tightly connected with follicular development [23]. During the post-emergence development in mosquitoes, JH drives sequential waves of gene expression [24]. Metabolic genes involved in glycolysis, glycogen/sugar metabolism, as well as the citrate cycle, are considerably repressed by JH in the previtellogenic phase, consistent with the accumulation of glycogen and triacylglycerols (TAG) in the sugar-fed mosquitoes [25]. These energy reserves are used later in vitellogenesis and their buildup in the previtellogenic phase determines the final quantity of eggs produced after a blood meal [26].

JH exerts its functions through the intracellular receptor Methoprene-tolerant (Met) [27], which belongs to the basic helix-loop-helix (bHLH)-Per-Arnt-Sim (PAS) transcription factor family [28]. Upon JH binding, Met forms a complex with Taiman (Tai), a transcriptional regulator that is essential for both JH and 20E responses [29–31]. The Met-Tai complex binds to the JH response elements (JHRE) in the JH-regulated genes and modulates their transcription [30]. JH induces rapid expression of several transcription factors, including Hairy and Krüppel homolog 1 (Kr-h1). These proteins act downstream from Met and regulate gene expression in response to JH [32]. On the other hand, 20E acts via a heterodimer of the ecdysone receptor (EcR) and Ultraspiracle (USP) to activate the expression of several transcription factors, including E74, E75, and HR3, which govern the expression of *vitellogenin* (*Vg*) and the development of follicles [32].

PPF is believed to affect mosquito reproduction by disturbing the normal action of endogenous hormones. Many studies of other insects have demonstrated that the anti-metamorphic action of JH analogs is mediated by Met [33,34]. An *in vitro* study has indicated that PPF binds *Tribolium castaneum* Met with higher affinity than JH-III [28]. Moreover, PPF has been shown to activate *Ae. aegypti* Met in cultured mosquito cells to regulate JH-inducible promoters [30,35]. While these lines of evidence suggest the potential involvement of Met in PPF-induced sterilizing effects in mosquitoes, this hypothesis has not been experimentally tested. Additionally, topical methoprene treatment has been reported to considerably decrease the expression of *USP*, *HR3*, and *Vg* at 18 h PBM in adult female *An. gambiae*, suggesting that methoprene causes a delay in vitellogenesis and egg maturation by repressing normal 20E response [18]. However, the 20E response in methoprene-treated mosquitoes has not been carefully investigated.

While many studies have confirmed the role of PPF in repressing mosquito reproduction [12,14–16], the mechanism underlying the sterilizing effect of PPF is poorly understood. In the current study, we initiated a comprehensive approach to systematically examine the physiological changes and developmental abnormalities invoked by PPF treatment. We monitored oocyte development and found that majority of primary follicles in PPF-treated mosquitoes were blocked in their development after blood feeding and failed to reach maturity. PPF caused metabolic shifting and disrupted normal hormone responses that are essential for egg maturation. These phenotypic changes correlated with the transcriptomic alterations revealed by RNA-seq analyses. Finally, we demonstrated that the JH receptor Met played a pivotal role in the PPF-caused reproductive disruption. These findings shed light on the molecular action of PPF in female sterilization and reveal potential molecular targets that could be exploited for the development of novel insecticides.

## Results

### PPF suppressed reproduction of female *Ae. aegypti* in a dose-dependent manner

The inhibitory effect of PPF on mosquito reproduction is affected by the time of exposure, formulation, methods of application, and other abiotic factors [36–38]. In this study, we exposed adult mosquitoes to PPF by placing them into cartons that had a lining of PPF-treated cotton gauzes. To establish a baseline for the sterilizing effect of PPF, adult female *Ae. aegypti* mosquitoes at 3 days after eclosion (PE) were exposed to increasing doses of PPF. Both untreated and cyclohexane-treated mosquitoes were used as control groups. Exposure to PPF considerably lowered the number of oviposited eggs and reduced hatching of eggs in a dose-dependent fashion (Table S1). A significant decrease in egg numbers was observed after exposure to PPF at 7 μg (active ingredient)/cm^2^ of the surface area for 30 min. At 70 μg/cm^2^, PPF caused a 94% reduction of egg numbers and prevented female mosquitoes from producing any viable offspring. Higher doses (e.g. 175 μg/cm^2^) appeared to be toxic to the exposed mosquitoes, decreasing mosquito survival and blood-feeding success (Table S1). In subsequent experiments of this study, mosquitoes were all exposed to PPF at 70 μg/cm^2^ for 30 min.

To examine whether sterilizing effect of PPF depends on the exposure time relative to blood-feeding, female adults from the same cohorts were treated with PPF (70 μg/cm^2^) for 30 min at 72 h PE, 96 h PE, 0.5 h PBM, 24 h PBM, or 36 h PBM (Fig S1). All the mosquitoes were given a blood meal at 120 h PE. PPF treatments in earlier stages of the first gonadotrophic cycle displayed more severe sterilizing effects than treatments in later stages. Exposure at 72 h PE exhibited the strongest impact, reduced egg production by 80%, and gave rise to no viable offspring (Table 1).

**Table 1:** Reproductive outcomes of five PPF treatment regimens.

| Regimens | Eggs/Female |  |  | Hatching rate |  |  | Relative reproduction rate |  |  |
| --- | --- | --- | --- | --- | --- | --- | --- | --- | --- |
|  | Unt | Cyc | PPF | Unt | Cyc | PPF | Unt | Cyc | PPF |
| **72 h PE** | 92.3<br>($\pm 4.2$) | 98.4<br>($\pm 3.1$) | 8.2<br>($\pm 5.9$)*** | 89.0<br>($\pm 4.6$) | 93.7<br>($\pm 0.6$) | 0.3<br>($\pm 0.6$)*** | - | 100 | 0.0<br>($\pm 0.0$)*** |
| **96 h PE** | 92.3<br>($\pm 4.2$) | 97.6<br>($\pm 10.0$) | 18.7<br>($\pm 3.6$)*** | 89.0<br>($\pm 4.6$) | 88.7<br>($\pm 1.5$) | 0.3<br>($\pm 0.6$)*** | - | 100 | 0.1<br>($\pm 0.1$)*** |
| **0.5 h PBM** | 92.3<br>($\pm 4.2$) | 82.5<br>($\pm 6.6$) | 34.9<br>($\pm 4.4$)*** | 89.0<br>($\pm 4.6$) | 89.0<br>($\pm 3.5$) | 3.3<br>($\pm 4.9$)*** | - | 100 | 1.5<br>($\pm 2.2$)*** |
| **24 h PBM** | 92.3<br>($\pm 4.2$) | 92.0<br>($\pm 2.8$) | 55.1<br>($\pm 3.2$)*** | 89.0<br>($\pm 4.6$) | 91.0<br>($\pm 2.6$) | 18.7<br>($\pm 12.1$)*** | - | 100 | 12.3<br>($\pm 8.3$)*** |
| **36 h PBM** | 92.3<br>($\pm 4.2$) | 91.1<br>($\pm 0.76$) | 66.6<br>($\pm 7.1$)** | 89.0<br>($\pm 4.6$) | 86.0<br>($\pm 7.0$) | 48.0<br>($\pm 6.6$)** | - | 100 | 40.7<br>($\pm 5.6$)*** |

Note: Unt, untreated; Cyc, cyclohexane-treated; PPF, pyriproxyfen-treated. \*, $p < 0.05$ ; \*\*, $p < 0.01$ ; \*\*\*, $p < 0.001$

The effects of PPF became weaker in other treatment regimens. Female mosquitoes exposed to PPF at 36 h PBM decreased their egg production and egg hatching by 27% and 44%, respectively, compared with the cyclohexane-treated group. Nevertheless, all five treatment regimens showed significant sterilizing effects. Egg retention increased substantially in PPF-treated mosquitoes, in accordance with the decline in egg oviposition (Fig S2). At 6 days after blood ingestion, 100% of female mosquitoes that were treated with PPF at 72 h PE or 96 h PE retained eggs, whereas egg retention was observed in 30% of females that were exposed to PPF at 36 h PBM (Fig S2).

After PPF exposure, female mosquitoes were dissected at 48 h PBM to examine the development of ovarian follicles. The development of primary follicles was severely impaired in mosquitoes that were treated with PPF at 72 h PE, 96 h PE, and 0.5 h PBM, as shown by a 47-54% decrease in follicular length (Table S2). The decrease also occurred, to a lesser extent, in mosquitoes exposed to PPF at 24 h PBM or 36 h PBM. Normal follicles at 48 h PBM had an elongated elliptical shape, whereas the follicles in PPF-treated mosquitoes were shorter in major axis and became spherical (Fig S3). Likewise, eggs laid by PPF-treated female mosquitoes displayed similar changes in egg shape as judged by the length-to-width ratios (Table S2, Fig S3).

### PPF exposure impaired ovarian follicular development

To characterize the effect of PPF on oogenesis, adult female mosquitoes (72 h PE) were exposed to PPF (70 μg/cm^2^) for 30 min. Follicular growth was compared between PPF-treated and cyclohexane-treated mosquitoes at multiple time points. Previtellogenic growth of primary follicles is normally completed by 72 h PE. Interestingly, PPF stimulated further growth of primary follicles before blood-feeding (Fig S4). At 120 h PE (48 h after PPF treatment), primary follicles in PPF-exposed mosquitoes were 40% longer in length than those of control mosquitoes. Primary follicles in control groups showed considerable growth in size after a blood meal; the average length of follicles increased from 131 μM (120 h PE) to 463 μM (48 h PBM). Conversely, the average follicular lengths in PPF-treated mosquitoes were 218 μM at 120 h PE and 221 μM at 48 h PBM (Fig S4), indicating that follicular growth in PPF-treated mosquitoes was nearly halted after blood feeding.

Next, we used Christopher’s stages of ovarian development [39,40] to define the effect of PPF on oocyte development. At 120 h PE, 79% of the primary follicles in control mosquitoes reached Christopher’s stage IIa while 20% were at stage IIb (Table 2). In contrast, 66% of those in PPF-treated females at the same time were at stage IIb and 32% at stage III, where the oocytes occupied at least 50% of the length of the follicle. Primary follicles in PPF-treated mosquitoes were generally larger at 120 h PE and loaded with more ooplasmic inclusions than those in control mosquitoes (Fig 1). At 24 h PBM, the vast majority of primary follicles in both PPF-treated and control groups developed to stage IIIb (>50-75% oocyte occupancy) (Table 2).

**Table 2.**
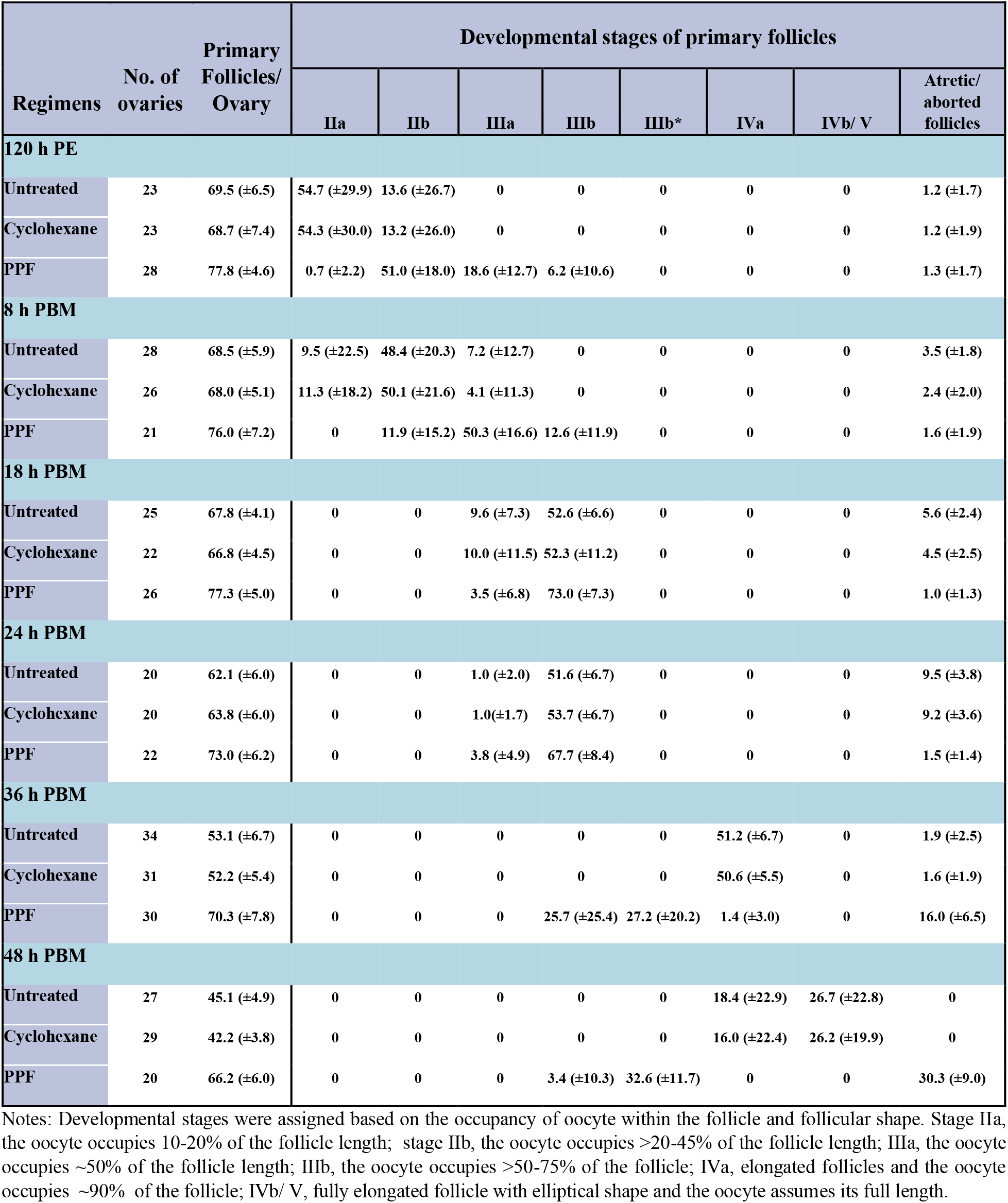
The Christopher’s stage of primary follicular development after PPF exposure.

**Fig 1.**
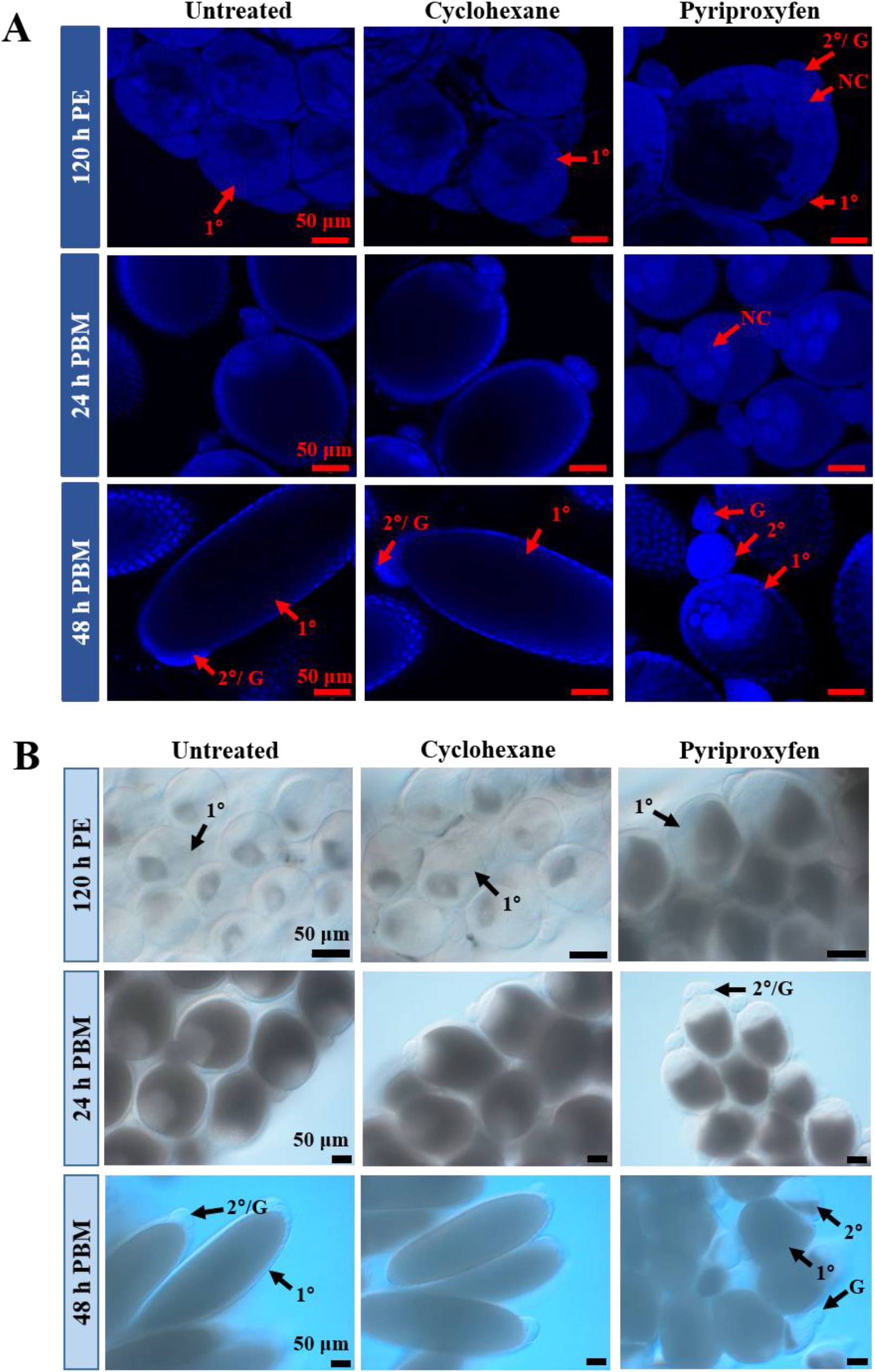
Follicular development in PPF-treated mosquitoes. (A) Ovarian follicles were collected at the indicated time points after PPF treatment. After fixation, follicles were stained with DAPI. Images were captured using a Zeiss LSM 880 confocal microscope. Scale bars represent 50 μm. (B) Differential interference contrast (DIC) images of ovarian follicles after PPF treatment. PE, Post eclosion; PBM, Post blood-meal. 1°, primary follicle; 2°, secondary follicle; G, germarium; NC, nurse cell.

However, follicles in PPF-treated mosquitoes were shorter in length and still occupied by large nurse cells (Fig 1A). About 14-15% of primary follicles in control groups were atretic/aborted, characterized by clearly visible small follicles and poor oocyte occupancy in the follicles; the PPF-treated counterparts had only 2% atretic/aborted follicles at this stage. Although not quantitatively measured, premature growth of secondary follicles was visibly prominent at 24 h PBM in PPF-treated mosquitoes, but not in control groups (Fig 1).

By 36 h PBM, around 96% of primary follicles in control mosquitoes entered stage IVa, where the follicles were slightly elongated, and the oocyte occupied ~90% of the space in the follicle. In PPF-treated mosquitoes, 37% of primary follicles remained at stage IIIb and 39% of the follicles were in an unusual stage (IIIb*), where the oocytes occupied ~90% of the follicles but kept the spherical shape (Table 2). Remarkably, 23% of the primary follicles in PPF-exposed mosquitoes were degenerated/aborted at 36 h PBM whereas only 3-4% follicles were atretic in control mosquitoes (Table 2).

A more striking difference was observed at 48 h PBM. More than 60% of primary follicles in control mosquitoes reached stage IVb/V, where nurse cells degenerated and disappeared (Table 2, Fig 1). In contrast, none of the primary follicles in PPF-treated mosquitoes entered stage IV. About 46% of them were arrested/aborted with clear signs of abortion (presence of nurse cells, disoriented vitelline ooplasm, and more opaque in appearance) (Fig S5), while another 49% of those follicles were stalled at stage IIIb*. Moreover, secondary follicles in PPF-treated mosquitoes showed additional growth and were loaded with ooplasmic inclusions (Fig 1B). Interestingly, although PPF treatment hindered oocyte development, it induced a significant increase in the number of primary follicles. Compared with the control groups, PPF-treated mosquitoes on average had 13% more primary follicles at 120 h PE and 57% more at 48 h PBM (Table 2).

### The abundances of storage macromolecules and circulating sugars were modified after PPF exposure

Nutritional status is an important indicator of reproductive fitness in any organism. To investigate whether PPF affects intermediary metabolism and energy storage, glycogen and triglyceride (TAG) were quantified in the whole body after female *Ae. aegypti* mosquitoes (72 h PE) were exposed to PPF. At 120 h PE, the glycogen levels in PPF-treated mosquitoes were 23% higher (*p* < 0.01) than those in cyclohexane-treated counterparts (Fig 2A). However, PPF seemed to have an opposite effect at 24 h PBM and caused a substantial depletion of glycogen, compared with the control mosquitoes. The measurements using colorimetric assays were validated by Periodic acid/Schiff (PAS) staining of mosquito fat bodies at those two time points (Fig 2B). PPF also displayed a similar effect on TAG levels in adult female mosquitoes. The amounts of TAG in PPF-treated mosquitoes were 34% higher than the cyclohexane control at 120 h PE but were 13% lower than the control groups at 24 h PBM (Fig 2C). The PPF-triggered alteration of TAG levels was substantiated by Nile red fluorescent staining of lipid droplets in the mosquito fat body (Fig 2D).

**Fig 2.**
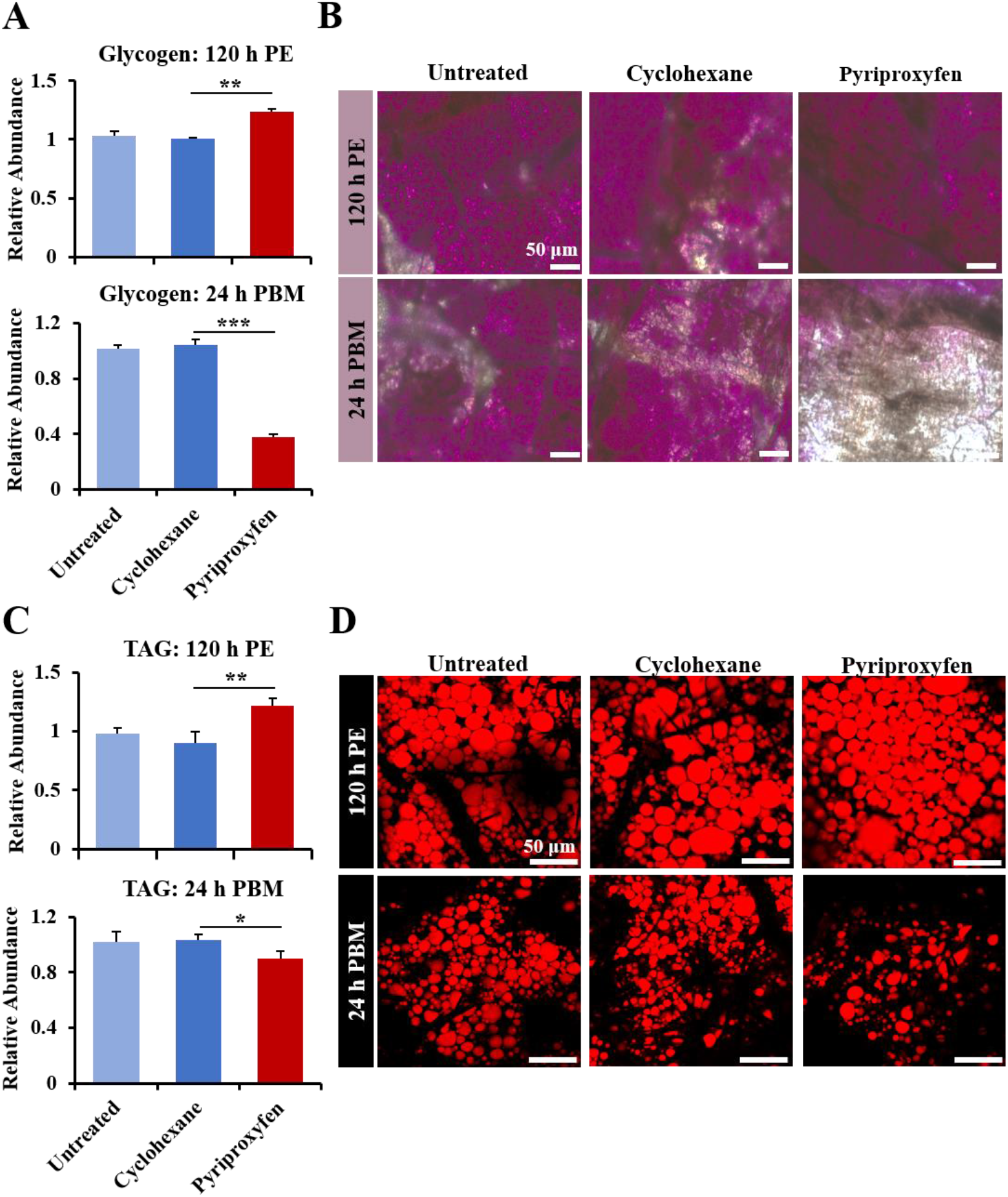
PPF exposure affected the levels of Glycogen and TAG. Female mosquitoes were treated with PPF (70 μg/cm^2^) at 72 h PE. (A) Quantitative measurement of glycogen in the whole-body mosquitoes at 120 h PE and 24 h PBM. The levels of glycogen were normalized to the protein contents of the samples. The relative abundance of the untreated control is set at 1. Error bars represent the standard deviation (SD) of three biological replicates. The differences between the PPF-treated and cyclohexane-treated mosquitoes were analyzed using paired t-test (*, *p* < 0.05; **, *p* < 0.01; ***, *p* < 0.001). (B) Glycogen in the fat body was visualized after PAS staining. The intensity of the pink/ purple color reflects the levels of glycogen. (C) Quantitative measurement of TAG in the whole-body mosquitoes. The levels of TAG were normalized to the protein contents of the samples. (D) Nile red lipid staining of lipid droplets in the fat body. Scale bar, 50 μm. PE, Post eclosion; PBM, Post blood-meal.

Additionally, the levels of four important circulating sugars (trehalose, sucrose, glucose, and fructose) were measured in the whole body of the mosquitoes. Compared with the solvent control, PPF treatment caused significant increases (*p* < 0.05) in the levels of sucrose, glucose, and fructose in the previtellogenic mosquitoes (120 h PE) (Table S3). At 24 h PBM, only sucrose was significantly more abundant (*p* < 0.01) in PPF-treated mosquitoes than the control groups. Thus, these data collectively indicated that PPF considerably altered energy mobilization and the metabolism of carbohydrates and lipids.

### The expression of metabolic genes was altered in the fat body after PPF treatment

To gain insights into PPF-induced female sterilization, global gene expression profiling was conducted using RNA-seq analyses. Female *Ae. aegypti* mosquitoes (72 h PE) were treated with PPF or cyclohexane. Both fat body and ovary tissues were collected after the treatment at 120 h PE and 24 h PBM. Dramatic changes in the transcriptomes were observed in the fat body as a result of PPF exposure. At 120 h PE, 1,457 *Ae. aegypti* genes were differentially expressed (log_2_|fold change| ≥ 0.8 and *padj* ≤ 0.01) in the fat body, with 558 genes upregulated and 899 genes downregulated in PPF-treated mosquitoes (Table S4). At 24 h PBM, 1,379 and 1,090 genes were upregulated and downregulated, respectively, in the fat body of PPF-treated mosquitoes when compared with the solvent control (Table S5). The differentially expressed genes belonged to a wide array of functional categories (Fig S6).

The fat body, which is functionally analogous to the vertebrate liver and adipose tissue, acts as a metabolic center and a storage organ in the mosquito [25,41]. Our transcriptome analysis revealed that 23% and 12.8% of differentially expressed genes at 120 h PE and 24 h PBM, respectively, were involved in the metabolism of carbohydrates and lipids (Tables S6-S9). For instance, the α-glucosidase 1 gene (*AAEL000223*), which encodes a glycogen breakdown enzyme, was significantly repressed in PPF-treated mosquitoes at 120 h PE (Table S6). Conversely, transcripts of this gene at 24 h PBM were 2.3-fold more abundant in PPF-treated mosquitoes than in cyclohexane-treated counterparts (Table S7). PPF treatment also reduced the expression of *enolase* (*AAEL001668*) at 120 h PE and lowered the expression of *hexokinase* (*HEX, AAEL009387*), *phosphoglycerate kinase* (*PGK*, *AAEL004988*), and *pyruvate kinase* (*PYK*, *AAEL012576*) at 24 h PBM (Tables S6 and S7). Weakened expression of the glycolytic enzymes may partly account for the increase in glucose in PPF-treated female mosquitoes. Moreover, several sugar transporters were differentially expressed in the fat body in response to PPF. The sucrose transporter gene (*STP 2*, *AAEL011520*) was activated in PPF-treated mosquitoes at 120 h PE but was downregulated at 24 h PBM (Tables S6 and S7). Another sucrose transporter gene (*STP 1*, *AAEL011519*) was repressed at both time intervals. Additionally, gene expression of nine sugar transporters was significantly altered at both 120 h PE and 24 h PBM after PFF exposure (Tables S6 and S7). Differential expression of *α-glucosidase 1, enolase, PGK, HEX* and *STP 2* in the fat body of PPF-treated mosquitoes was validated using real-time PCR (Fig 3).

**Fig 3.**
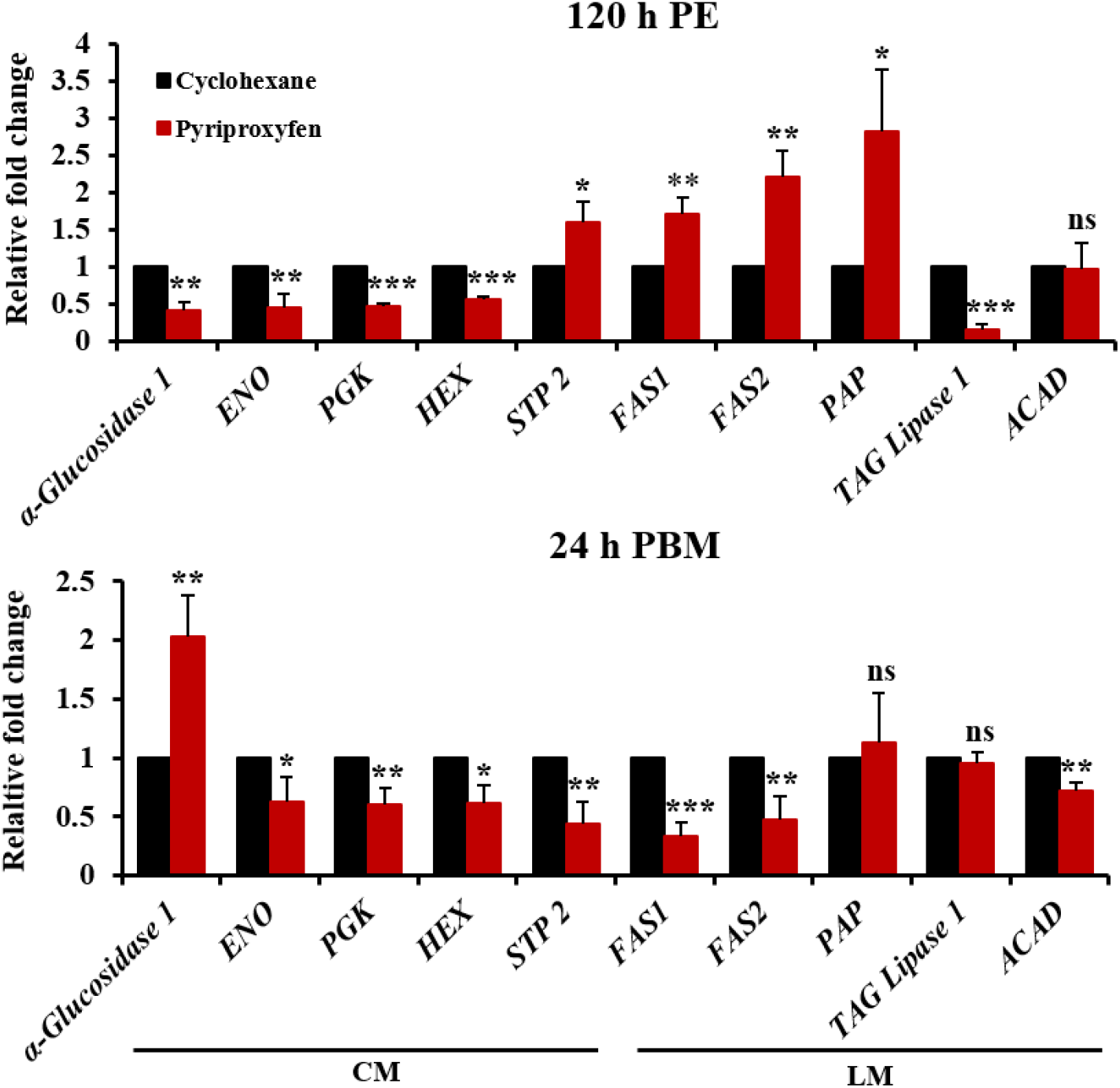
PPF treatment altered the expression of metabolic genes. Female mosquitoes were treated with PPF at 72 h PE. Quantitative RT-PCR assays were performed to compare the expression of selected genes involved in carbohydrate metabolism (CM) and lipid metabolism (LM). mRNA levels were measured in the fat body at 120 h PE and 24 h PBM. Data are presented as mean ± SD from three independent replicates. Statistical analyses were performed using paired t-test (ns, *p* > 0.05; *, *p* < 0.05; **, *p* < 0.01; ***, *p* < 0.001).

Lipid metabolic pathway was also affected by PPF treatment. At 120 h PE, the expression of several lipid synthesis enzymes, including fatty acid synthase 1 (FAS1, *AAEL008160*), fatty acid synthase 2 (FAS2, *AAEL002228*), and phosphatidate phosphatase (PAP), was considerably enhanced in PPF-treated mosquitoes (Table S8, Fig 3). In contrast, several enzymes that are involved in lipid degradation, such as TAG lipase 1, were downregulated by PPF at 120 h PE. The differential expression seemed to correlate well with the elevated accumulation of lipid droplets in the fat body at 120 h PE after PPF exposure (Fig 2). The effect of PPF on lipid metabolic pathway was more complicated after blood-feeding. The expression of *FAS1 and FAS2* was repressed at 24 h PBM after PPF treatment (Table S9, Fig 3). Besides, PPF exposure led to lower expression at 24 h PBM of Acyl-CoA dehydrogenase (ACAD) (Fig 3), which catalyzes the initial step of fatty acid beta-oxidation. In mosquitoes, storage lipids are crucial for egg maturation and act as a fuel for ~90% of the energy required during embryogenesis [42]. Several proteins that play important roles in lipid transportation were downregulated in the fat body after PPF exposure. The expression of two lipid transport genes (monocarboxylate transporter, *AAEL002412,* and *AAEL008347*) was significantly reduced at 24 h PBM in PPF-treated mosquitoes, compared with the cyclohexane-treated group (Table S5). Likewise, PPF exposure reduced the expression of *apolipophorin III* (*AAEL008789*) at 24 h PBM in the fat body.

PPF also affected other important functions of the fat body. ABC transporters are involved in the translocation of lipids, sugars, and amino acids in insects [43]. Three ABC transporter genes (*AAEL008672*, *AAEL014699*, *AAEL008138*) were significantly repressed at 24 h PBM in PPF-exposed mosquitoes (Table S5). Additionally, programmed autophagy in the mosquito fat body is essential for the normal progression of gonadotropic cycles [41]. Two autophagy-related genes, *APG8* (*AAEL007162*) and *APG4B* (*AAEL007228*) were significantly suppressed at 24 h PBM after PPF exposure at 72 h PE (Table S5), potentially contributing to the aberrant follicular development after PPF treatment.

### PPF caused transcriptomic changes in the ovary

RNA-seq analyses of the ovary tissues revealed 862 and 653 differentially expressed genes at 120 h PE and 24 h PBM, respectively, in PPF-treated mosquitoes (Tables S10 and S11). Some of the differentially expressed genes are known for their roles in follicular development. Vitelline membrane proteins are released from follicular epithelium after blood-feeding and are involved in the formation of the innermost layer of eggshell [44]. Vitelline membrane proteins genes *15a-1* (*AAEL013027*) and *15a-3* (*AAEL014561*) displayed 49-fold and 172-fold upregulation, respectively, at 120 h PE in the ovaries of PPF-treated mosquitoes, compared with the control mosquitoes (Table S10). However, the expression of the two genes was considerably lower in PPF-treated mosquitoes than cyclohexane-treated ones at 24 h PBM (Table S11). Cytoskeleton proteins are required for maintaining the shape of follicle and egg via cytoskeletal reorganization [45]. *Calponin* (*AAEL008303*) and *myosin motor protein* (*AAEL012543*) were repressed at 24 h PBM in response to PPF exposure. The homeobox protein SIX4 (*AAEL010327*) regulates oocyte development in *Ae. aegypti*; its ectopic expression after blood feeding leads to smaller follicle size and significantly reduced egg numbers [46]. The mRNA levels of *SIX4* in PPF-treated mosquitoes were 2-fold higher than those in the control mosquitoes at 24 h PBM (Table S11). On the other hand, the PPF treatment caused a 4.4-fold reduction at 24 h PBM in the mRNA levels of FOXL (*AAEL005741*), a forkhead transcription factor that is essential for oocyte development and egg deposition in *Ae. aegypti* [47]. Overall, PPF modulated the expression of numerous genes that are associated with follicular development and egg maturation.

### PPF alters the expression patterns of 20E response genes

RNA-seq analysis also indicated that PPF treatment markedly altered the expression of several well-characterized 20E response genes both before and after blood-feeding. The differential expression between PPF-treated and control mosquitoes was validated by quantitative RT-PCR. At 120 h PE, at a time when 20E was at the background level, PPF significantly induced the expression of *EcRA*, *E75A*, *HR3*, *HR4*, and *Vg* in the fat body (Fig S7). The first 4 genes encode important transcriptional regulators in the 20E response. Vg is the major yolk protein precursor synthesized by mosquito fat body. In the ovary, the mRNA levels of *EcRA*, *EcRB*, *E75A*, *HR3*, and *HR4* at 120 h PE were all considerably higher in PPF-treated mosquitoes than control mosquitoes (Fig S7). When the 20E titers peaked at around 24 h PBM, PPF treatment seemed to have the opposite effect. These 20E response genes were all expressed, in the fat body and ovary, at significantly lower levels in PPF-treated mosquitoes than cyclohexane-treated counterparts (Fig S7). In contrast, *Krüppel homolog 1* (*Kr-h1*), a JH-inducible gene, showed 2.2-fold upregulation at 120 h PE in the fat body and ovary after PPF exposure. The PPF-induced expression of *Kr-h1* was even more robust at 24 h PBM (Fig S7).

The expression of these 20E response genes was further compared between PPF-treated and control mosquitoes in the fat body throughout the first gonotrophic cycle. The mRNA levels of *EcRA* and *E75A* were significantly enhanced at 96 and 120 h PE in PPF-treated mosquitoes (Fig 4). They lacked the 20E-induced elevation at 16, 24, and 30 h PBM as displayed in the control mosquitoes but showed a considerable increase later at 36 h PBM. The expression of *HR3* and *Vg* was induced in response to PPF in the previtellogenic stage and exhibited further increase after blood-feeding. However, the expression of *HR3* and *Vg* after a blood meal was considerably weakened in PPF-treated mosquitoes, compared with the control group (Fig 4). On the other hand, significant overexpression of *Kr-h1* was detected from 120 h PE to 24 h PBM after PPF exposure. Therefore, the results indicated that PPF treatment considerably elevated expression of Kr-h1 and altered the expression patterns of 20E response genes during egg maturation.

**Fig 4.**
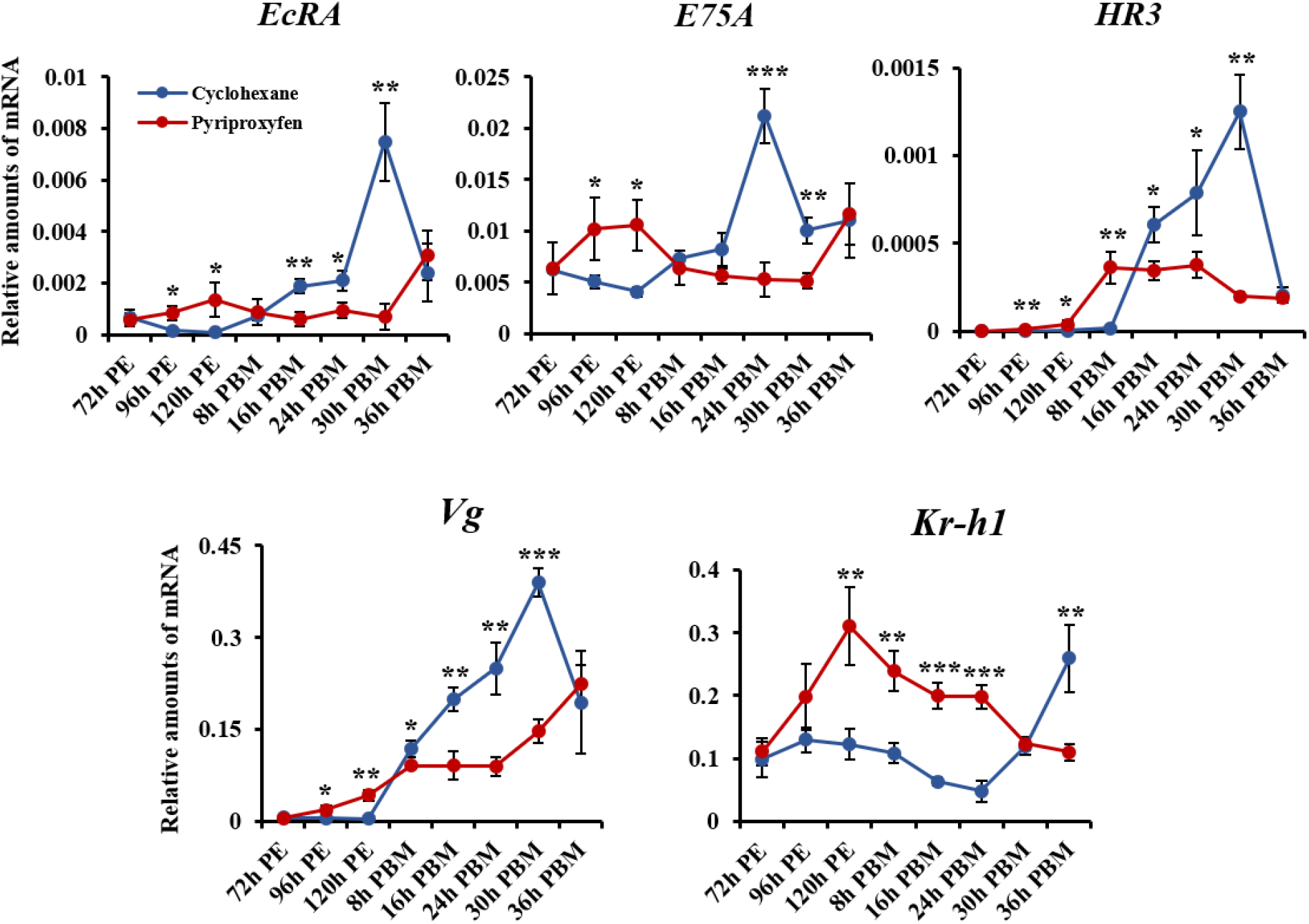
Expression patterns of selected hormone-regulated genes in PPF-treated mosquitoes. The mRNA levels were analyzed in the fat body of female mosquitoes at the indicated time points after PPF exposure at 72 h PE. The cyclohexane-treated mosquitoes were used as a control group. Transcript abundance was normalized to that of the Ribosomal Protein S7 (RpS7) gene. Data are shown as the mean ± standard deviation of three biological replicates. Statistical analysis was performed using a Student’s t-test (ns, *p* > 0.05; *, *p* < 0.05; **, *p* < 0.01; ***, *p* < 0.001).

### PPF-induced sterilization effect was mediated through the JH receptor Met

To determine the role of Met in the PPF-induced sterilization, we examined how the Met-dependent genes in the previtellogenic stage responded to PPF treatment. A previous study reported that the expression of 2,554 genes was significantly altered after RNAi knockdown of *Met* (iMet) in *Ae. aegypti* fat body during the previtellogenic stage [48]. Among those Met-dependent genes, 345 genes showed significant changes (|log2 fold change| ≥ 0.8, *padj* ≤ 0.05) in their mRNA levels in PPF-treated mosquitoes, compared with cyclohexane-treated counterparts (Fig S8).

To further explore the role of Met in PPF-treated mosquitoes, RNAi-mediated knockdown of Met was performed in adult female *Ae. aegypti* mosquitoes. The mRNA levels of *Met* were considerably reduced at 120 h PE and 24 h PBM in female mosquitoes injected with double-stranded RNA (dsRNA) for *Met*, compared with the cohorts that were injected with control dsRNA (Fig 5A and 5B). Injection of dsMet substantially reduced the expression of selected 20E responsive genes (*Vg*, *HR3*, *E75A*, and *EcRA*) at 120 h PE in both PPF-treated and cyclohexane-treated mosquitoes. Notably, the PPF-induced overexpression of these genes at 120 h PE was abrogated in the Met-depleted mosquitoes (Fig 5A). Knockdown of Met lowered the expression of *Vg*, *HR3*, *E75A,* and *EcRA* at 24 h PBM in PPF-treated and control mosquitoes (Fig 5B). The depletion of Met could not rescue the repression of these genes by PPF at 24 h PBM. The expression of two carbohydrate metabolic genes (*sucrose transporter protein STP 2* and *α-glucosidase 1*) and two lipid metabolic genes (*phosphatidic acid phosphatase/* (*PAP*) and *TAG lipase 1*) were also analyzed in *Met* RNAi mosquitoes. At 120 h PE, the mRNA levels of *STP 2* in the PPF-treated and cyclohexane-treated mosquitoes were not significantly affected by the knockdown of *Met* (Fig S9). In contrast, injection of dsMet caused a marked increase in the mRNA abundance of *α-glucosidase 1* and *TAG lipase 1* in the PPF-treated and cyclohexane-treated mosquitoes; the PPF-caused downregulation of both genes was abolished in the *Met* RNAi mosquitoes (Fig S9). These results together indicated that Met plays an important role in the PPF-altered expression of 20E response genes and selected metabolic genes, especially in the previtellogenic phase.

**Fig 5.**
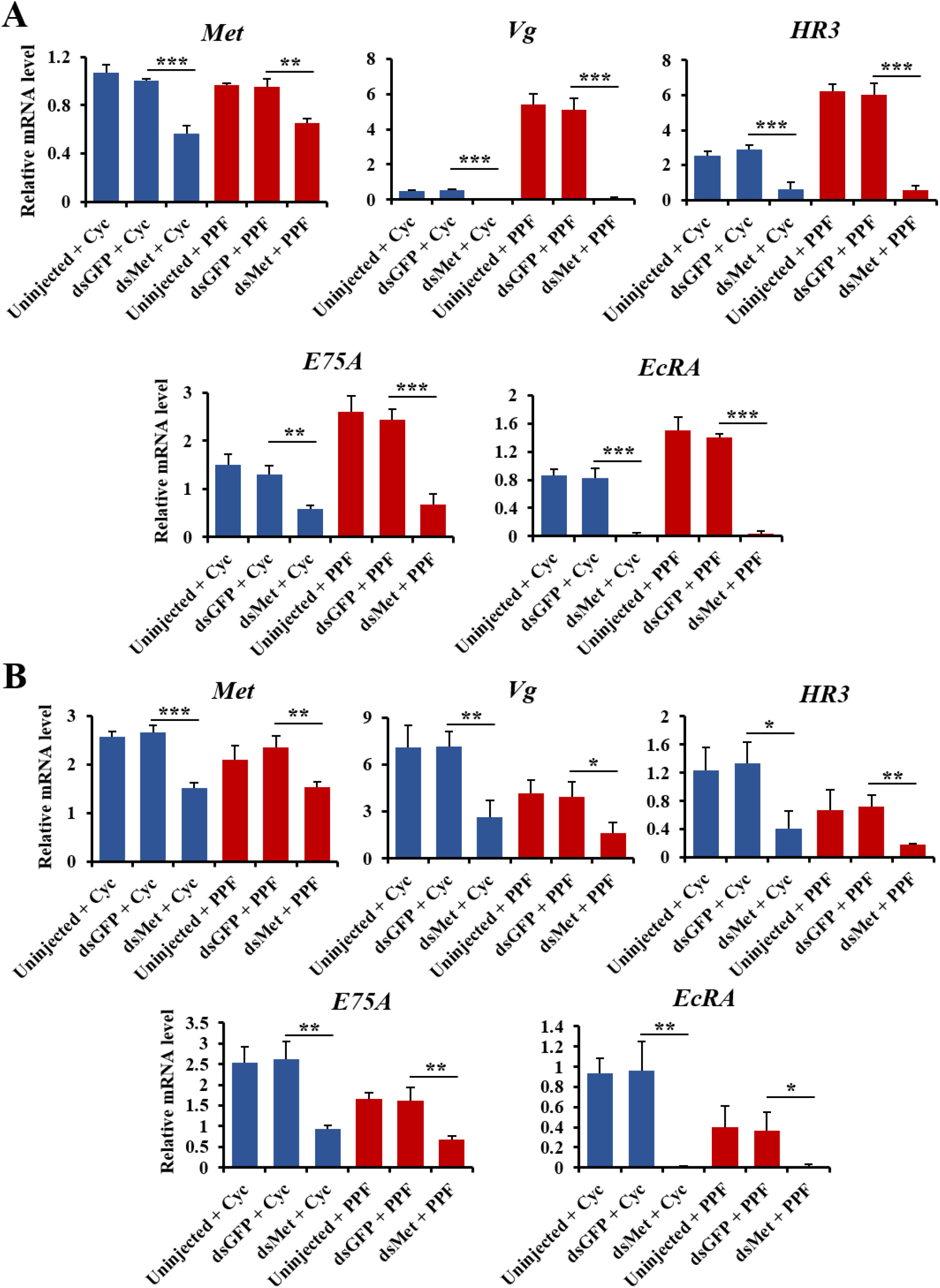
Effect of Met depletion on the PPF-triggered alterations of gene expression. DsMet was injected into adult female mosquitoes within 1 h after eclosion. Untreated and dsGFP-injected mosquitoes were used as control groups. At 72 h PE, the *Met* RNAi mosquitoes and control groups were treated with cyclohexane (Cyc) or PPF (70 μg/cm^2^). Fat bodies were collected at 96 h PE (A) or 24 h PBM (B). Relative mRNA levels of the indicated genes were determined using real-time PCR. Error bars represent the standard deviation of three replicates. Statistical significance was determined using a Student’s t-test (ns, *p* > 0.05; *, *p* < 0.05; **, *p* < 0.01; ***, *p* < 0.001).

Follicular development was assessed after PPF exposure in Met-depleted adult female mosquitoes. The post-eclosion growth of primary follicles, normally controlled by endogenous JH, was severely impaired in the dsMet-injected mosquitoes as expected (Fig 6A). The PPF-induced overgrowth in the previtellogenic stage (120 h PE) was also abrogated by the depletion of Met, which caused a reduction of length by 62% compared to the PPF-treated control mosquitoes (Fig 6A and 6E). Although the growth of primary follicles was still repressed by PPF treatment after blood feeding, the length of primary follicles in the Met-deficient mosquitoes was 12% longer at 48 h PBM than the dsGFP-injected and PPF-treated counterpart (Fig 6A). Moreover, the PPF-triggered premature growth of secondary follicles was blocked after blood-feeding in the dsMet-injected female mosquitoes, but not in the PPF-treated control groups (Fig 6E).

**Fig 6.**
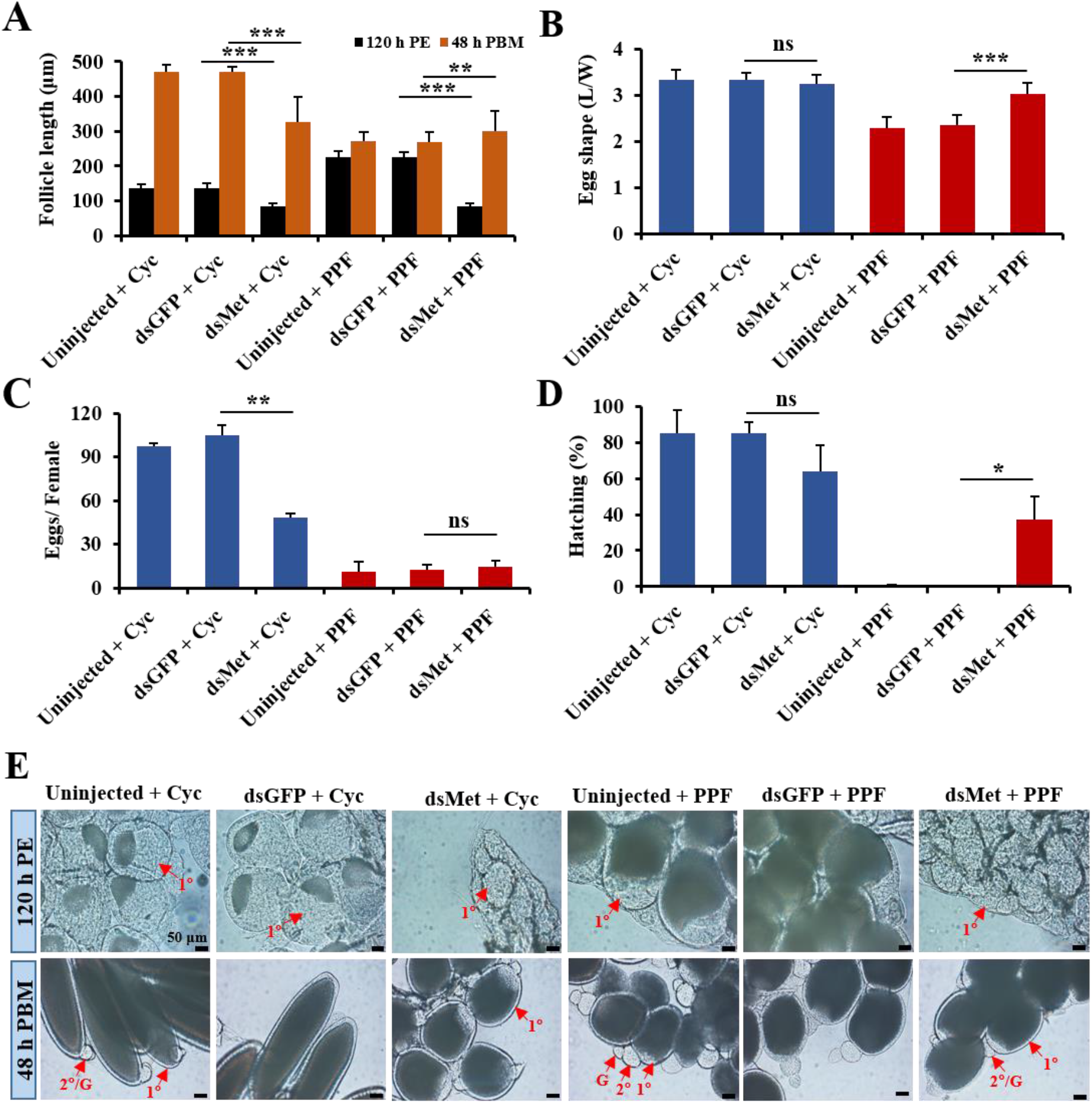
Implication of Met in the impaired egg maturation in the PPF-treated female mosquitoes. Adult female mosquitoes were injected with dsRNAs within 1 h after eclosion. The injected and uninjected mosquitoes were treated with cyclohexane (Cyc) or PPF at 72 h PE. All the mosquitoes were given a blood meal at 120 h PE. (A) Length of primary follicles. Follicular lengths (N=30 follicles) were measures at 120 h PE and 48 h PBM. (B) Egg shape parameter (egg L/W ratio) was calculated after measuring the lengths and widths of 30 eggs. (C) Average number of eggs produced by each female mosquito. Oviposited eggs were counted at 6 days PBM. Data are shown as the mean ± standard deviation of three biological replicates (10 mosquitoes/replicate). (D) Hatching rates for different treatment groups. Eggs were allowed to hatch for 7 days to record the hatching rate. Error bars illustrate the standard deviation of three biological replicates (10 mosquitoes/replicate). Statistical analysis was conducted using paired t-test (ns, *p* > 0.05; *, *p* < 0.05; **, *p* < 0.01; ***, *p* < 0.001). (E) Ovarian follicular development. Follicles were dissected at 120 h PE and 48 h PBM. Differential Interference Contrast (DIC) images were taken using Zeiss Axiocam ERc5. PE: Post eclosion, PBM: Post blood-meal, 1°: primary follicle, 2°: secondary follicle, G: germarium.

The knockdown of Met did not significantly increase the number of eggs produced by the PPF-treated female mosquitoes after a blood meal (Fig 6B). However, the eggs laid by the dsMet-injected females after PPF treatment had an elongated shape more resembling normal eggs; the egg shape parameter (L/W ratio of laid eggs) of the PPF-treated mosquitoes was increased by 29% in the dsMet-injected females, compared with the dsGFP-injected group (Fig 6C). Most importantly, depletion of Met partially rescued poor hatching of the eggs laid by the PPF-treated females; the hatching rate went up from 0% for the dsGFP-injected group to 37% for the dsMet-injected group (Fig 6D). In summary, Met plays a pivotal role in every aspect of the sterilizing effect of PPF in *Ae. aegypti* female mosquitoes. It is required for the PPF-caused alteration of 20E response, hampering follicle development, and impairment of ultimate reproductive success.

## Discussion

The objective of this study was to elucidate the mechanistic details in the PPF-caused sterilization of female mosquitoes. Our results demonstrate that PPF was highly effective to reduce the reproductive output of female *Ae. aegypti*. The development of primary follicles in PPF-treated mosquitoes was severely compromised while the secondary follicles were prematurely activated. The impaired follicular development was likely caused by a disruption of normal genetic programs that are coordinated by insect hormones and also by alterations of metabolism and energy mobilization after PPF treatment. This was substantiated by our transcriptomic analysis which revealed massive changes in the expression patterns of many genes related to 20E hormonal cascades and various metabolism. Additionally, we have found that depletion of the JH intracellular receptor Met ameliorated several anomalies resulted from the PPF treatment and partially rescued the infertility of PPF-treated mosquitoes, therefore establishing a pivotal role of Met in mediating the sterilization effect of PPF.

We have shown that PPF exposure in the early stages of the first gonadotrophic cycle had more detrimental effects on mosquito reproduction than exposure in the late stages. This is in agreement with an early study reporting that the fecundity of the PPF-treated *Ae. aegypti* females decreased further with increasing time between PPF treatment and blood-feeding [13]. Similar observations were also reported with *An. arabiensis, An. gambiae, Culex quinquefasciatus* mosquito species [15,16,37]. This conserved phenomenon in different mosquito species suggests that PPF influences multiple steps in egg maturation and that the final reproductive outcome of PPF-treated female mosquitoes reflects the cumulative effects of PPF.

Alternating waves of two major insect hormones, JH and 20E, regulate follicular development and reproductive maturation in mosquitoes. Rising JH titers after eclosion stimulate the growth of primary follicles. After blood ingestion, levels of JH drop rapidly to near background levels while 20E titers increase and peak at 18-24 h post a blood meal. The uptake of PPF, a JH mimic, is believed to perturb the normal hormonal profiles in adult female mosquitoes. JH-regulated growth of primary follicles was normally completed by 72 h PE before the PPF treatment was given in our study. However, PPF seemed to stimulate the deposition of more vitelline granules and yolk proteins into ovarian follicles between 72 h PE and blood-feeding. As a result, the follicles of PPF-treated mosquitoes proceeded to more advanced developmental stages, compared with control mosquitoes. While JH is thought to have no role in the stimulation of protein synthesis in the mosquito fat body [49], we found the JH analog PPF significantly increased the mRNA levels of *Vg* in the fat body of PPF-exposed female mosquitoes during the previtellogenic stage. Overexpression of *Vg* after PPF exposure could account for the increased sizes of the primary follicles in the PPF-treated mosquitoes at 120 h PE.

The growth of secondary follicles is normally controlled by JH that starts to rise at the end of vitellogenesis. Although the maturation of primary follicles was stalled in the PPF-treated mosquitoes after blood feeding, the secondary follicles manifested premature growth at 48 h PBM. The activation of secondary follicles is usually suppressed by oostatic hormone before the primary oocyte has reached maturity [50,51]. It will be interesting to investigate whether PPF suppresses the action of oostatic hormone and initiates the development of secondary follicles despite incomplete maturation of primary follicles.

Follicular resorption/degeneration is common during the maturation of follicles [52]. Matured follicles observed after blood-feeding are fewer than the primary follicles present after eclosion. We observed a significant increase in the number of primary follicles in PPF-treated females throughout the first gonotrophic cycle, which could result from decreased resorption of follicles after PPF treatment. At 36 and 48 h PBM, PPF-treated mosquitoes were loaded with a significant number of aborted follicles, which were characterized by the dark color, relatively small sizes, and diffused/disorganized vitellus granules. Higher numbers of primary follicles have been previously reported in *Ae. aegypti* and *Cx. pipiens* (L.) females after topical application of methoprene [53]. A very small portion of the primary follicles reached maturity in the PPF-treated mosquitoes. The poor hatching success of the few eggs laid by PPF-treated females is likely due to an aberrant embryonic development, which exhibits disrupted differentiation and no segmentation [12].

The reproductive output of a female mosquito is dependent on nutrient resources and regulated by hormonal responses [54]. Carbohydrate and lipid metabolic pathways in mosquitoes are directly regulated by JH and 20E, leading to the accumulation of glycogen and TAG during the previtellogenic stage and the decline of these energy reserves after blood ingestion [25][55]. Therefore, the JH analog PPF may affect metabolic homeostasis through metabolic reprogramming. Indeed, PPF treatment further enhanced the buildup of glycogen and TAG in the previtellogenic female adults and accelerated the depletion of these storage macromolecules after a blood meal. Transcriptome analysis of PPF-treated mosquitoes substantiated the metabolism shifting. Several enzymes (α-glucosidase, lipase, TAG lipase, etc.) that are involved in glycogen and lipid breakdown were significantly repressed by PPF in the previtellogenic stage but were upregulated after blood-feeding. Anabolic enzymes, such as GLY, FAS-1, 2, etc., also exhibited altered expression patterns in the PPF-treated mosquitoes, corroborating the PPF-enhanced glycogen and TAG levels at 120 h. Through modulating the expression of metabolic genes, PPF could disrupt many physiological processes that are essential for egg development and maturation. For example, lipids are crucial energy storage that makes up a large part of insect yolk. Lipid mobilization from the fat body to ovaries increases after blood-feeding to support follicular development [56]. Thereby, the decrease of TAG in PPF-treated mosquitoes could potentially reduce lipid deposition in the developing oocytes and impair egg maturation.

PPF treatment also significantly altered the levels of sucrose, glucose, and fructose but not trehalose in the exposed mosquitoes. It is unknown whether changes in these important circulating sugars contribute to the compromised reproduction of PPF-treated female adults. The metabolic shifting after PPF treatment may be a component of stress responses [57,58]. After PPF exposure, *Bombyx mori* also increases the levels of intermediary metabolites, including glucose [59].

An important function of JH in the previtellogenic adult female mosquitoes is to prepare mosquito tissues for the stage-specific response to 20E [60,61]. Topical application of methoprene, another JH analog, represses the expression of 20E responsive genes during the vitellogenic stage in *An. gambiae* [18]. Our study indicated that PPF modulated 20E-regulated genes in both the previtellogenic and vitellogenic phases. PPF considerably enhanced the expression of *EcRA*, *E75A*, *HR3*, and *Vg* at 120 h PE, at a time when 20E remained at the low basal levels. After a blood meal, PPF imposed the opposite effect and substantially dampened the 20E-induced expression of these genes. In female mosquitoes, 20E plays a central role in regulating vitellogenesis and egg development. The aberrant expression patterns of these 20E-regulated transcriptional factors in PPF-treated mosquitoes undoubtedly contributed to the disrupted egg maturation. The mechanism underlying the PPF-modulated expression of 20E response genes is largely unknown. PPF considerably enhanced the expression of Kr-h1, a JH-regulated transcriptional factor, both before and after blood feeding. Kr-h1 was first identified as a stage-specific modulator of the prepupal ecdysone response in *Drosophila melanogaster* [61]. In the early immature stages of insects, Kr-h1 prevents precocious metamorphosis by directly repressing 20E-dependent activation of the pupal specifier gene *BR-C* and the adult specifier gene *E93* [62,63]. Therefore, the elevated expression of Kr-h1 in PPF-treated female mosquitoes might directly or indirectly modify the expression of the aforementioned 20E regulated genes. More experiments are needed to test this hypothesis.

Many studies have shown that PPF is an agonist ligand of the JH receptor Met and the action of PPF is mediated by Met in insects [28,33]. Our transcriptomic analysis confirmed that a considerable portion of Met-dependent genes displayed significant differential expression after PPF exposure. Moreover, using selected 20E response genes and metabolic genes, we demonstrated that depletion of Met abrogated many of the PPF-induced changes in mRNA abundance. Knockdown of Met in the previtellogenic female mosquitoes hampered the follicular development and altered the expression of 20E responsive gene expression in untreated and cyclohexane-treated mosquitoes. In PPF-treated female adults, the PPF-induced previtellogenic overgrowth of primary follicles and the premature activation of secondary follicles vanished after the expression of Met in adult mosquitoes was knocked down using RNAi. Although the depletion of Met did not increase the numbers of oviposited eggs, it drastically boosted the hatching rate of the eggs laid by PPF-treated females. Therefore, these results collectively indicated that Met is an important mediator of the sterilizing effect of PPF in mosquitoes. Met is indispensable for the JH-regulated post-eclosion development in female mosquitoes [30,48]. The knockdown of Met in the previtellogenic stage makes it difficult to separate the effects of PPF from those of endogenous JH. A possible solution is to inject dsRNA for *Met* at 72 h PE, at a time when the post-eclosion development is completed. The injected mosquitoes will then be exposed to PPF at 0.5 h PBM as PPF treatment after a blood meal still substantially reduced mosquito fecundity (Table 1). It is important to note that among the PPF-regulated genes in the previtellogenic mosquitoes, only a small portion of them (~24%) was found to be Met-dependent. This suggests that Met is not the only molecular target of PPF in mosquitoes and other players are involved in mediating the action of PPF in mosquito reproduction. The Met-independent PPF action should be further elucidated in future studies.

## Methods

### Mosquito rearing

The *Ae. aegypti* Liverpool strain was reared as described previously [64]. Mosquitoes were maintained at 28°C and 60–70% relative humidity on a 14:10 h light: dark cycle. Larvae were reared in big plastic pans and fed with pulverized fish food (Tetra, Blacksburg, VA) and protein pellets. Newly emerged adults were placed in collapsible cages (12 × 12 × 12 in, BioQuip Products, Rancho Dominguez, CA) and provided with a 10% (wt/vol) sucrose solution daily using cotton pads. To initiate vitellogenesis and egg production, female mosquitoes (5 days post eclosion) were fed on defibrinated sheep blood (Colorado Serum Company, Denver, CO) using an artificial membrane feeder.

### Pyriproxyfen treatment

Pyriproxyfen (NyGuard®; MGK, Minneapolis, MN) was dissolved in cyclohexane at various concentrations. Gauze pads (62 cm × 18.5 cm; Equate), which were to simulate a bed net, were soaked in PPF solutions for 1 hour and then allowed to air-dry completely. Gauzes treated with either PPF or cyclohexane were used as a liner inside mosquito cartons (16 oz. double poly-coated paper cups). Mosquito groups (~40 females/ replicate) from the same cohort were anesthetized by short exposure to cold and transferred to the abovementioned cartons. After 30 min of confinement, mosquitoes were returned to new untreated cartons for subsequent assays. All experiments were performed in triplicate, each being repeated at least three times with different batches of mosquitoes.

### Assessment of mosquito fecundity and fertility

After PPF treatment, mortality was recorded at 24 h post treatment and dead mosquitoes were discarded. Female mosquitoes were blood-fed at 120 h post eclosion, regardless of whether the mosquitoes were exposed to PPF before or after a blood meal. Fully engorged mosquitoes were kept for subsequent experiments. Each replicate contained ~40 females. Ovaries were dissected at 48 h PBM to examine oocyte development. The length of primary ovarian follicles was measured using Leica Application Suite (v4.5). For egg collection, ten blood-fed female mosquitoes from each replicate were transferred into a new cup and oviposition sites were set up at 48 h PBM. Egg papers were collected at 144 h PBM and eggs were counted manually. To determine the hatch rate, ~100 eggs were randomly selected from each replicate and transferred into separate plastic cups filled with water to allow them to hatch. The number of mosquito larvae was recorded for 7 days. All experiments were repeated at least three times with different cohorts of mosquitoes. The relative reproductive rate was calculated for each PPF concentration, compared with cyclohexane control. Relative reproduction rate = the average number of larvae produced by each PPF-treated female/the average number of larvae produced by each cyclohexane-treated female.

### Measurements of glycogen and triglyceride

Glycogen levels were determined colorimetrically at 540 nm using the Glycogen assay kit (Sigma-Aldrich) following the manufacturer’s instructions. Three independent biological samples were used for each experimental condition and each sample contained the whole body of six adult female mosquitoes. The results were normalized to the protein contents in the sample, which were quantified using BCA protein assay (Thermo Fisher Scientific). Periodic Acid-Schiff (PAS) staining was used to visualize glycogen in the fat body of female mosquitoes following the protocol of Yamada et al. [65].

TAG was measured in mosquito whole body extracts as described previously [25]. A colorimetric assay was performed using the Triglyceride Colorimetric Assay Kit (Cayman) following the manufacturer’s instructions. TAG contents were normalized to the protein levels of the samples. Nile Red fluorescent dye was used to stain lipid droplets in the fat body as reported by Wang et al. [66].

### High-performance thin-layer chromatography (HPTLC)

To quantify trehalose, sucrose, glucose, and fructose after PPF exposure, HPTLC was performed as described by Fell et al. [67]. Briefly, 25 mosquitoes from each experimental group were homogenized with 70% ethanol. After centrifugation at 13000 × g for 5 min, the supernatant was spotted onto pre-coated HPTLC silica gel plates (10 cm × 10 cm). Quantitative measurements were made by analyzing the sugar spots with ImageJ software (NIH, MD).

### Examination of follicular development

Dissected ovaries were treated with PBS-T (Phosphate buffered saline with 0.5% Triton X-100) for 5 minutes at room temperature to increase the permeability of follicle cells. Subsequently, ovarian follicles were incubated with DAPI (4′,6-diamidino-2-phenylindole, 100 nM) for 30 seconds in a dark place and imaging was performed using a Zeiss 880 confocal microscope. DAPI staining was observed at excitation/emission: 358 nm/461 nm. Zeiss Axiocam ERc5 was used for differential interference contrast (DIC) imaging. Christopher’s stages of ovarian development were used to define follicular development in the first gonotrophic cycle [39,40].

### RNA-seq and data analysis

Transcriptomes in the fat body and ovary of the PPF-treated female mosquitoes were analyzed using RNA-seq. Cyclohexane-treated mosquitoes were included as the control group. Three biological replicates were taken, and each replicate contained fat bodies or ovaries isolated from 10 mosquitoes. Total RNA was extracted using TRIzol reagent (Invitrogen) and the Direct-zol miniprep plus kit (Zymo Research). mRNAs were then purified from the total RNA using NEBNext Poly(A) mRNA Magnetic Isolation Module (New England Biolabs). mRNA samples were sent to the Novogene corporation (Sacramento, CA, USA), where RNA-seq libraries were prepared and sequenced using an Illumina platform (paired-end 150 bp reads).

Sequence reads were aligned using TopHat (v2.1.1) [68] and bowtie (v2.2.5) [69] to the *Ae. aegypti* reference genome (AaegL3) [70]. HTSeq-count [71] was then used to calculate the number of reads for each gene and mRNA abundance was normalized based on FPKM values (Fragments Per Kilobase of transcript per Million mapped reads). Gene expression was compared between the PPF-treated and cyclohexane-treated mosquitoes using the DESeq package in R [72]. Differential expression was defined by ≥1.75- fold change and an adjusted false-discovery rate (*Padj* value) of ≤ 0.01 [48]. Functions of differentially expressed genes were predicted using EggNOG (evolutionary genealogy of genes: Non-supervised Orthologous Groups, v5.0) [73]. RNA-seq data have been submitted to the NCBI Sequence Read Archive (SRA) repository (accession number: PRJNA599428).

### Quantitative Real-Time PCR

Each sample contained fat bodies or ovaries dissected from 10 female mosquitoes. Total RNA was extracted using TRIzol reagent (Invitrogen) and the Direct-zol miniprep plus kit. cDNA synthesis was performed from 1 μg of total RNA using the Omniscript RT Kit (Qiagen). All the samples were analyzed in triplicate using GoTaq qPCR Master Mix (Promega) on an ABI 7300 system (Applied Biosystems). Gene-specific primers are listed in Table S12. Transcript abundance was normalized to the expression of the *RPS7* gene (*AAEL009496*).

### RNAi-mediated gene silencing

Expression of the *Met* gene was knocked down by injection of double-stranded RNA (dsRNA) into newly emerged adult female mosquitoes as described previously [64]. DsRNA for the enhanced green fluorescent protein (EGFP) was used as a control. Within 1 h of adult emergence, 1 μg of dsMet was injected into each female mosquito. At 72 h PE, both control groups (untreated and dsGFP-injected) and the dsMet-injected group were treated with cyclohexane solvent or PPF (70 μg/cm^2^ for 30 min). Fat body samples were collected at 96 h PE and 24 h PBM to evaluate the knockdown efficiency of *Met*.

## Supporting information

Schematic diagram of PPF exposure at various stages in adult female mosquitoes

Egg retention after different PPF treatments

Effect of PPF exposure on the morphology of follicles and eggs

Growth of primary follicles after PPF exposure

PPF hampered follicular development of mosquitoes

Differential gene expression in the fat body and ovary after PPF treatment

Expression of 20E response genes in PPF-treated mosquitoes

Differential expression of Met-dependent genes in response to PPF treatment

PPF-triggered alterations of gene expression in the Met-depleted female mosquitoes

Dose-dependent effect of PPF on Ae. aegypti reproduction

Effect of PPF exposure on follicle/egg morphology

Amounts of circulating sugars after PPF exposure

List of DEGs in the fat body at 120 h PE transcriptome

List of DEGs in the Fat body at 24 h PBM transcriptome

list of carbohydrate pathway related gene expression level in the fat body at 120 h PE transcriptome

List of carbohydrate pathway related gene expression level in the fat body at 24 h PBM transcriptome

Lipid synthesis and breakdown related gene expression level in the fat body at 120 h PE transcriptome

Lipid synthesis and breakdown related gene expression level in the fat body of female Ae. aegypti at 24 h PBM transcriptome

DEGs in the female Ae. aegypti ovary at 120 h PE transcriptome

DEGs in the Ae. aegypti ovary at 24 h PBM transcriptome

List of primers used in quantitative real-time PCR and dsRNA synthesis for RNAi mediated knockdown

## Abbreviations

20E: 20 hydroxyecdysone
EcR: Ecdysone receptor
JH: Juvenile hormone
Kr-h1: Krüppel homolog 1
Met: Methoprene-tolerant
PPF: Pyriproxyfen
PE: Post-eclosion
PBM: Post blood-meal
TAG: Triacylglyceride
USP: Ultraspiracle
Vg: Vitellogenin

## Acknowledgments

The authors would like to thank Drs. Aaron Gross, Chloé Lahondère, and Clément Vinauger for their helpful advice and constructive suggestions, and thank Ms. Sandra Gabbart for assisting us with the high-performance thin-layer chromatography experiment.

## Supporting Information

### Supplementary figure captions

**Fig. S1. Schematic diagram of PPF exposure at various stages in adult female mosquitoes.** After adult emergence, both male and female *Ae. aegypti* mosquitoes were allowed to mate for three days. Female mosquitoes were treated with PPF (70 μg/cm^2^) for 30 minutes at the indicated time points. All the mosquitoes were given a blood meal at 120 h after eclosion. Follicle examination was performed at 48 h PBM. Oviposited eggs were counted six days after blood feeding. Matured eggs were allowed to hatch for seven days to record the hatching rate. PE, Post eclosion; PBM, Post blood-meal.

**Fig. S2. Egg retention after different PPF treatments.** Adult female mosquitoes were exposed to PPF at the indicated time points. Mosquitoes from untreated, cyclohexane-treated, and PFF-treated groups were dissected six days after blood-feeding. Developed follicles inside the abdomen after oviposition were counted as ‘retained eggs’. Reported here are the average of retained eggs in each female mosquito (10 mosquitoes/ replicate) and the percentages of females with retained eggs (10 mosquitoes/ replicate). Results shown are the mean ± S.D. of three replicates. Statistical differences between the PPF-treated and cyclohexane-treated mosquitoes were analyzed using paired t-test (ns, *p* > 0.05; *, *p* < 0.05; **, *p* < 0.01; ***, *p* < 0.001).

**Fig. S3. Effect of PPF exposure on the morphology of follicles and eggs.** Female adults were exposed to PPF (70 μg/cm^2^) at 72 h PE and given a blood meal at 120 h PE. Thirty mosquitoes, from untreated, cyclohexane-treated, and PPF-treated groups, were dissected at 48 h PBM to examine ovarian follicles. Eggs were collected at 144 h PBM and imaging was performed using Leica Application Suite (v4.5). Scale bars represent 0.1 mm in the upper and lower panels. PBM, Post blood-meal.

**Fig. S4. Growth of primary follicles after PPF exposure.** Adult female mosquitoes were treated with PPF at 72 h PE. The lengths of primary follicles were measured at the indicated time points throughout the first gonotrophic cycle. Error bars represent the standard deviation (SD) of three biological replicates. Statistical analysis was performed by paired t-test. Asterisks indicate significant differences in follicular length between cyclohexane-treated and PPF-treated groups (ns, *p* > 0.05; *, *p* < 0.05; **, *p* < 0.01; ***, *p* < 0.001). PE, Post-eclosion; PBM, Post blood-meal.

**Fig. S5. PPF hampered follicular development of mosquitoes.** PPF treatment was carried out at 72 h PE in *Ae. aegypti* female mosquitoes. Shown here with an asterisk sign (*) is a representative image of aborted ovarian follicles in PPF-treated mosquitoes at 48 h PBM. These follicles are characterized by their round shape, advance development of secondary follicle/prominent germarium, and the presence of intact nurse cells, which normally disappear in control mosquitoes at this stage. 1°, primary follicle; 2°, secondary follicle; G, germarium; NC, nurse cell.

**Fig. S6. Differential gene expression in the fat body and ovary after PPF treatment.** Adult female mosquitoes were treated with PPF (70 μg/cm^2^) at 72 h PE. Cyclohexane was used as a solvent control. RNA-seq analyses were performed using mosquito fat body and ovary tissues collected at 120 h PE and 24 h PBM. The percentage of differentially expressed genes in discrete functional categories is displayed in the pie chart. PE, Post eclosion; PBM, Post blood-meal.

**Fig. S7. Expression of 20E response genes in PPF-treated mosquitoes.** Adult female mosquitoes were treated with PPF at 72 h PE. The expression of selected 20E-and JH-response genes was analyzed using real-time PCR. Fold changes of mRNA abundance in the fat body or ovary of PPF-treated mosquitoes were determined relative to the cyclohexane-treated control group. Data are presented as mean ± SD from three independent replicates. Statistical analysis was performed using paired *t*-test (ns, *p* > 0.05; *, *p* < 0.05; **, *p* < 0.01; ***, *p* < 0.001).

**Fig. S8. Differential expression of Met-dependent genes in response to PPF treatment.** (A) An overlap between genes that were modulated by PPF treatment and those that were affected by depletion of Met (iMet) in the previtellogenic fat body. The Met-dependent genes were reported by Zou et al. [54]. (B) Scatter plot of the 345 PPF genes that were significantly altered in response to both PPF treatment and RNAi-mediated knockdown of Met. Log2 fold changes (lfc) of iMet (Met RNAi vs control RNAi) were shown in X axis, whereas log2 fold changes after PPF exposure (PPF treatment vs control treatment) were represented in Y axis. Each dot illustrates a single gene.

**Fig. S9. PPF-triggered alterations of gene expression in the Met-depleted female mosquitoes.** DsMet was injected into adult female mosquitoes within 1 h after eclosion. Untreated and dsGFP-injected mosquitoes were used as control groups. At 72 h PE, the *Met* RNAi mosquitoes and control groups were treated with cyclohexane (Cyc) or PPF. Fat bodies were collected at 96 h PE (A) or 24 h PBM (B). Relative mRNA levels of indicated genes were determined using real-time PCR. Error bars represent the standard deviation of three replicates. Statistical significance was conducted using a Student’s t-test (ns, *p* > 0.05; *, *p* < 0.05; **, *p* < 0.01; ***, *p* < 0.001).

Table S1. Dose-dependent effect of PPF on *Ae. aegypti* reproduction

Table S2. Effect of PPF exposure on follicle/egg morphology

Table S3. Amounts of circulating sugars after PPF exposure

Table S4. Differentially expressed genes in the fat body at 120 h PE after PPF treatment

Table S5. Differentially expressed genes in the fat body at 24 h PBM after PPF treatment

Table S6. PPF-induced differential expression of carbohydrate metabolic genes in the fat body at 120 h PE

Table S7. PPF-induced differential expression of carbohydrate metabolic genes in the fat body at 24 h PBM

Table S8. PPF-induced differential expression of lipid metabolic genes in the fat body at 120 h PE

Table S9. PPF-induced differential expression of lipid metabolic genes in the fat body at 24 h PBM

Table S10. PPF-induced differential gene expression in the ovary at 120 h PE

Table S11. PPF-induced differential gene expression in the ovary at 24 h PBM

Table S12. List of primers used in quantitative real-time PCR and dsRNA synthesis

