## Supplementary figures and images for "The juvenile hormone receptor Methoprene-tolerant is involved in the sterilizing effect of pyriproxyfen on adult *Aedes aegypti* mosquitoes"

### Differential expression of Met-dependent genes in response to PPF treatment

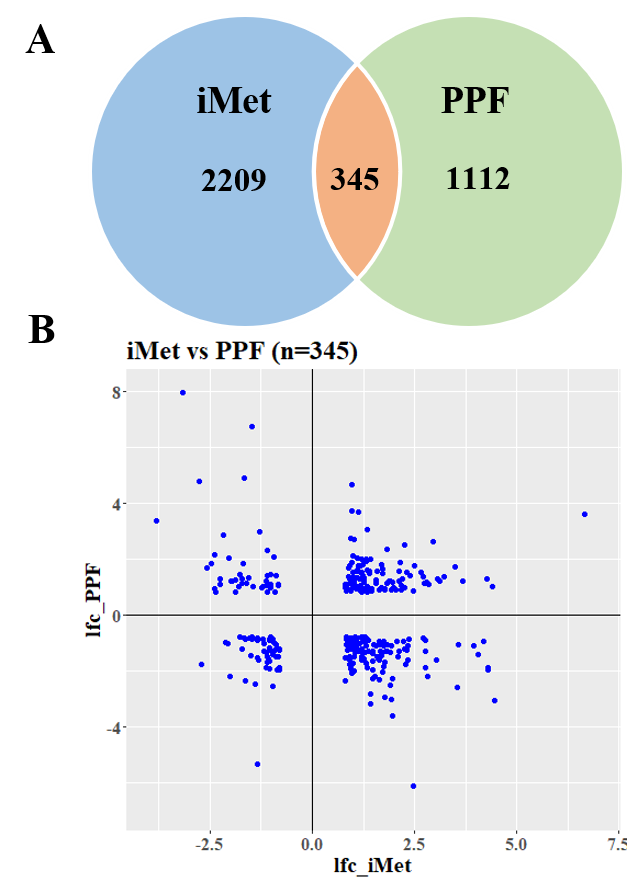

### Differential gene expression in the fat body and ovary after PPF treatment

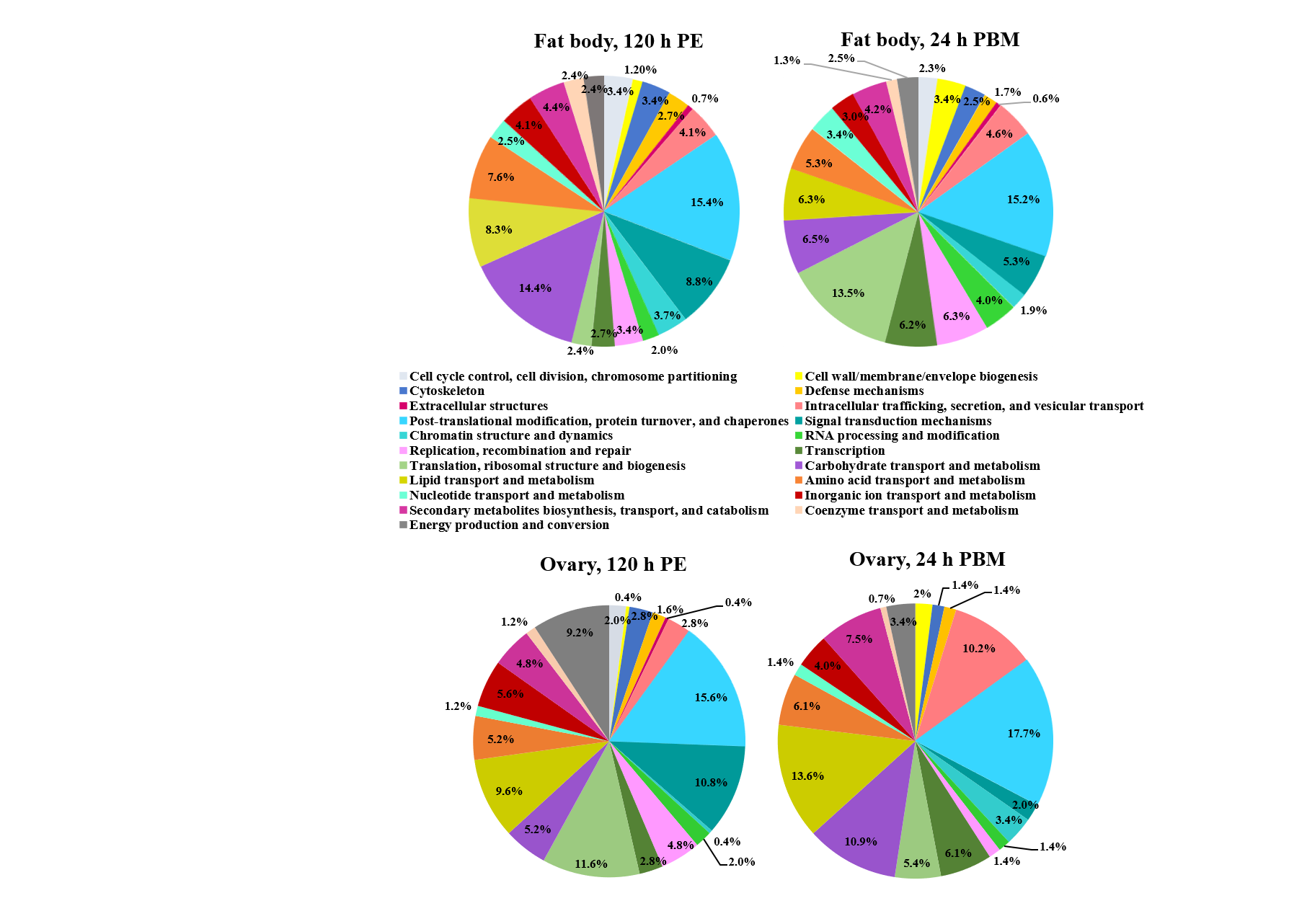

### Effect of PPF exposure on the morphology of follicles and eggs

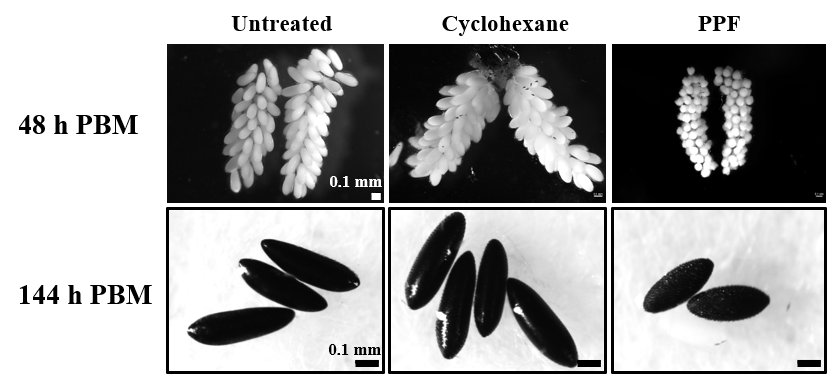

### Egg retention after different PPF treatments

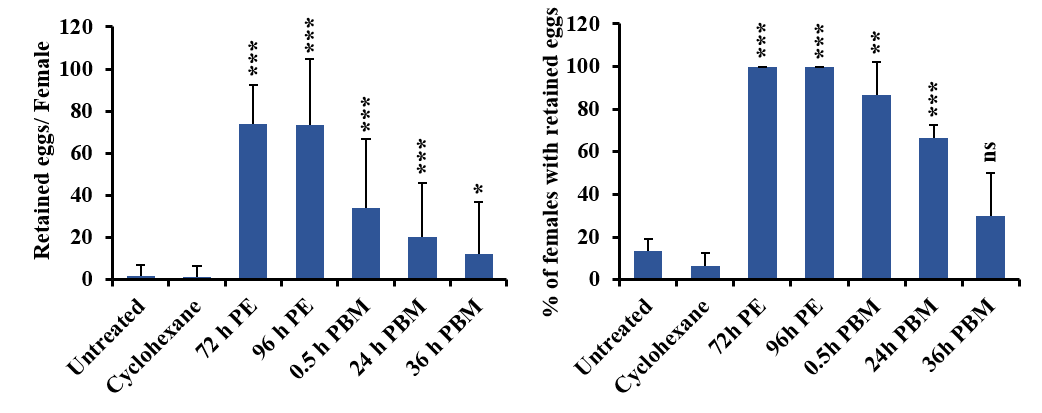

### Expression of 20E response genes in PPF-treated mosquitoes

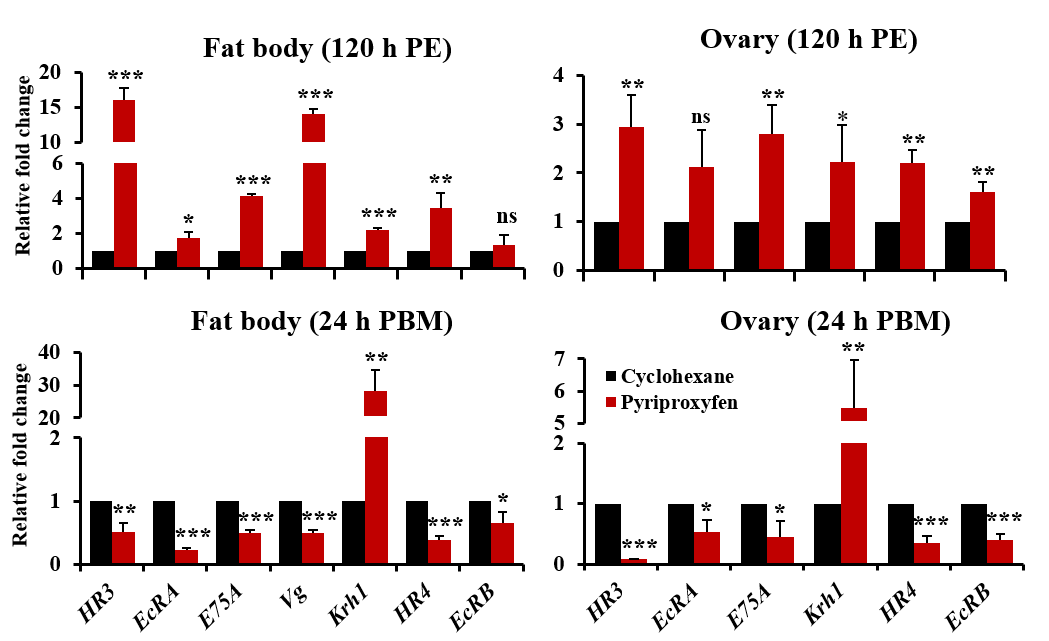

### Growth of primary follicles after PPF exposure

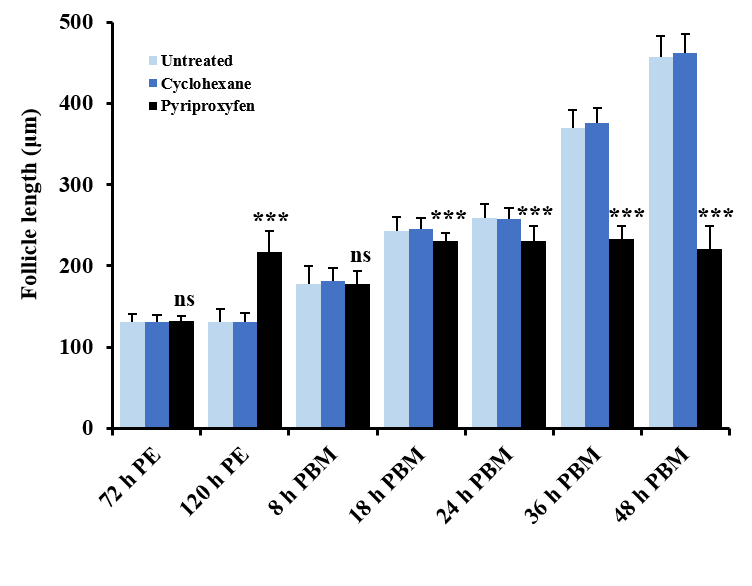

### PPF hampered follicular development of mosquitoes

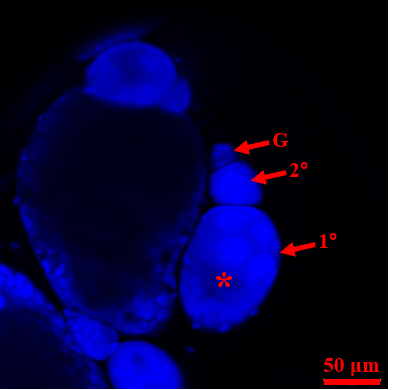

### PPF-triggered alterations of gene expression in the Met-depleted female mosquitoes

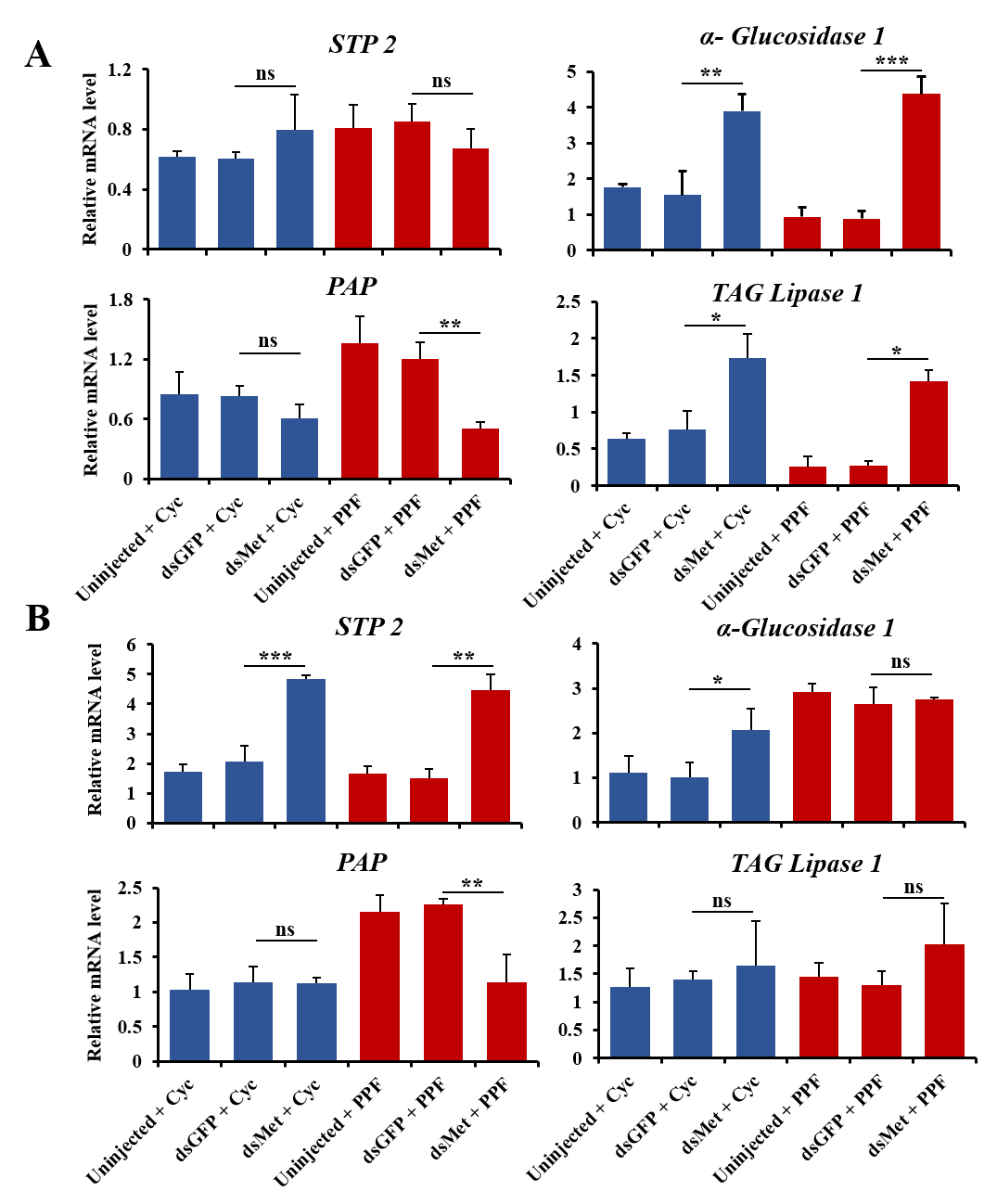

### Schematic diagram of PPF exposure at various stages in adult female mosquitoes

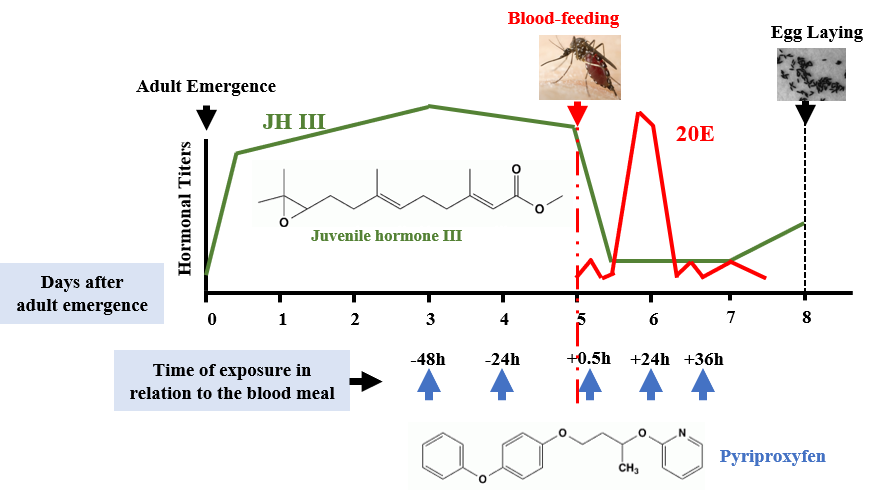
