## Supplementary material for "The juvenile hormone receptor Methoprene-tolerant is involved in the sterilizing effect of pyriproxyfen on adult *Aedes aegypti* mosquitoes": Dose-dependent effect of PPF on Ae. aegypti reproduction

**Table S1. Dose-dependent effect of PPF on *Ae. aegypti* reproduction**

| Groups | % Mortality | % Blood-fed | Eggs/Female | Hatching rate | Relative reproduction rate |
| --- | --- | --- | --- | --- | --- |
| Untreated | 0.0 (±0.0) | 98.2 (±1.5) | 90.0 (±4.6) | 88.0 (±1.0) | - |
| Cyclohexane | 0.0 (±0.0) | 98.0 (±1.8) | 91.6 (±9.5) | 84.0 (±1.7) | 100 |
| PPF 3.5 µg/cm^2^ | 0.9 (±1.6) ^ns^ | 94.3 (±5.3)^ns^ | 90.2 (±5.8)^ns^ | 83.0 (±6.2)^ns^ | 97.3 (±9.5)^ns^ |
| PPF 7 µg/cm^2^ | 1.0 (±1.7) ^ns^ | 92.4 (±1.3)^*^ | 56.9 (±6.6)^**^ | 71.3 (±5.5)^*^ | 52.5 (±2.3)^***^ |
| PPF 10.5 µg/cm^2^ | 1.0 (±1.7) ^ns^ | 91.4 (±2.5)^*^ | 56.9 (±9.3)^*^ | 33.7 (±3.8)^***^ | 24.7 (±2.4)^***^ |
| PPF 14 µg/cm^2^ | 1.1 (±1.9) ^ns^ | 92.0 (±2.4)^*^ | 58.7 (±6.0)^**^ | 26.7 (±4.5)^***^ | 20.2 (±2.5)^***^ |
| PPF 17.5 µg/cm^2^ | 1.8 (±0.3) ^ns^ | 92.1 (±1.1)^*^ | 48.8 (±8.2)^**^ | 24.7 (±5.0)^***^ | 16.0 (±5.9)^***^ |
| PPF 26.3 µg/cm^2^ | 1.8 (±1.6) ^ns^ | 92.5 (±1.6)^*^ | 42.0 (±8.2)^**^ | 9.9 (±9.5)^***^ | 4.8 (±6.0)^***^ |
| PPF 35 µg/cm^2^ | 1.8 (±1.6) ^ns^ | 92.1 (±2.5)^*^ | 20.3 (±10.6) ^***^ | 5.7 (±2.3)^***^ | 1.4 (±0.9)^***^ |
| PPF 70 µg/cm^2^ | 1.8 (±1.6) ^ns^ | 92.0 (±3.0)^*^ | 5.7 (±3.7)^***^ | 0.0 (±0.0)^***^ | 0.0 (±0.0)^***^ |
| PPF 105 µg/cm^2^ | 7.0 (±3.5)^*^ | 92.3 (±1.5)^*^ | 7.4 (±7.5)^***^ | 0.0 (±0.0)^***^ | 0.0 (±0.0)^***^ |
| PPF 175 µg/cm^2^ | 39.1 (±11.4)^**^ | 82.0 (±2.7)^**^ | 4.7 (±4.0)^**^ | 0.0 (±0.0)^***^ | 0.0 (±0.0)^***^ |

Note: Mortality was measured at 24 h after PPF exposure. After blood-feeding, only fully engorged mosquitoes from each experimental group were kept for the measurements of eggs/female, hatching rate and relative reproduction rate determination. ns, *p* > 0.05; *, *p* < 0.05; **, *p* < 0.01; ***, *p* < 0.001
