## Supplementary material for "The juvenile hormone receptor Methoprene-tolerant is involved in the sterilizing effect of pyriproxyfen on adult *Aedes aegypti* mosquitoes": Effect of PPF exposure on follicle/egg morphology

**Table S2. Effect of PPF exposure on follicle/egg morphology**

| **Regimens** | **Treatment** | **Follicle length (μm)** | **Egg length (L) (μm)** | **Egg width (W) (μm)** | **Egg shape (L/W)** |
| --- | --- | --- | --- | --- | --- |
|  | **Untreated** | 445 (±20.4) | 584.9 (±17.9) | 171.5 (±13.4) | 3.4 (±0.3) |
| **72 h PE** | **Cyclohexane** | 456.4 (±25.1) | 593.2 (±25.1) | 173.7 (±10.1) | 3.4 (±0.3) |
|  | **PPF** | 211.1 (±13.3)^***^ | 446.1 (±42.6)^***^ | 183.3 (±10.2)^***^ | 2.4 (±0.3)^***^ |
|  | **Untreated** | 445 (±20.4) | 584.9 (±17.9) | 171.5 (±13.4) | 3.4 (±0.3) |
| **96 h PE** | **Cyclohexane** | 449.3 (±25.6) | 586.9 (±28.7) | 169.5 (±8.5) | 3.5 (±0.2) |
|  | **PPF** | 234.7 (±14.5)^***^ | 457.7 (±29.6)^***^ | 173.6 (±14.4)^ns^ | 2.8 (±0.3)^***^ |
|  | **Untreated** | 445 (±20.4) | 584.9 (±17.9) | 171.5 (±13.4) | 3.4 (±0.3) |
| **0.5 h PBM** | **Cyclohexane** | 447.5 (±25.0) | 593.7 (±23.6) | 171.4 (±11.4) | 3.5 (±0.2) |
|  | **PPF** | 237.8 (±15.9)^***^ | 452.5 (±36.5)^***^ | 167.0 (±13.0)^ns^ | 2.6 (±0.4)^***^ |
|  | **Untreated** | 445 (±20.4) | 584.9 (±17.9) | 171.5 (±13.4) | 3.4 (±0.3) |
| **24 h PBM** | **Cyclohexane** | 449.7 (±23.4) | 583.5 (±28.6) | 171.5 (±9.5) | 3.4 (±0.2) |
|  | **PPF** | 351.3 (±31.2)^***^ | 535.5 (±33.9)^***^ | 169.2 (±10.6) ^ns^ | 3.2 (±0.3)^***^ |
|  | **Untreated** | 445 (±20.4) | 584.9 (±17.9) | 171.5 (±13.4) | 3.4 (±0.3) |
| **36 h PBM** | **Cyclohexane** | 452.1 (±23.1) | 595.7 (±19.8) | 168.8 (±5.9) | 3.5 (±0.1) |
|  | **PPF** | 430.4 (±48.7)^*^ | 541.0 (±37.6)^***^ | 172.2 (±8.2) ^ns^ | 3.1 (±0.2)^***^ |

Note: Statistical differences between the PPF-treated and cyclohexane-treated mosquitoes were analyzed using paired t-test. ns, *p* > 0.05; *, *p* < 0.05; **, *p* < 0.01; ***, *p* < 0.001
