## Supplementary material for "The juvenile hormone receptor Methoprene-tolerant is involved in the sterilizing effect of pyriproxyfen on adult *Aedes aegypti* mosquitoes": Amounts of circulating sugars after PPF exposure

**Table S3. Amounts of circulating sugars after PPF exposure**

| Groups | Trehalose  (μg/ female) | Sucrose  (μg/ female) | Glucose  (μg/ female) | Fructose  (μg/ female) |
| --- | --- | --- | --- | --- |
| **120 h PE** | | | | |
| **Untreated** | 5.41 (±0.60) | 21.24 (±0.03) | 11.36 (±0.44) | 5.16 (±0.27) |
| **Cyclohexane** | 5.01 (±0.45) | 21.37 (±0.27) | 11.22 (±0.37) | 5.48 (±0.16) |
| **PPF** | 4.09 (±0.81) **^ns^** | 23.17 (±0.29)^*^ | 13.07 (±0.36)^*^ | 7.54 (±0.12)^**^ |
| **24 h PBM** | | | | |
| **Untreated** | 3.61 (±0.21) | 16.07 (±0.81) | 6.0 (±0.31) | 5.50 (±0.10) |
| **Cyclohexane** | 3.51 (±0.16) | 16.11 (±0.57) | 6.06 (±0.35) | 5.72 (±0.07) |
| **PPF** | 3.50 (±0.08) **^ns^** | 20.96 (±0.33)^**^ | 7.44 (±0.77) **^ns^** | 6.73 (±1.06) **^ns^** |

Note: ns, *p* > 0.05; *, *p* < 0.05; **, *p* < 0.01
